# Hierarchical recurrent state space models reveal discrete and continuous dynamics of neural activity in *C. elegans*

**DOI:** 10.1101/621540

**Authors:** Scott Linderman, Annika Nichols, David Blei, Manuel Zimmer, Liam Paninski

## Abstract

Modern recording techniques enable large-scale measurements of neural activity in a variety of model organisms. The dynamics of neural activity shed light on how organisms process sensory information and generate motor behavior. Here, we study these dynamics using optical recordings of neural activity in the nematode *C. elegans*. To understand these data, we develop state space models that decompose neural time-series into segments with simple, linear dynamics. We incorporate these models into a hierarchical framework that combines partial recordings from many worms to learn shared structure, while still allowing for individual variability. This framework reveals latent states of population neural activity, along with the discrete behavioral states that govern dynamics in this state space. We find stochastic transition patterns between discrete states and see that transition probabilities are determined by both current brain activity and sensory cues. Our methods automatically recover transition times that closely match manual labels of different behaviors, such as forward crawling, reversals, and turns. Finally, the resulting model can simulate neural data, faithfully capturing salient patterns of whole brain dynamics seen in real data.

## Introduction

New recording technologies are transforming neuroscience, allowing us to measure neural activity, sensory stimuli, and natural behavior with unprecedented scale and precision. Our aim is to discover simplifying structure in these high-dimensional recordings, but the dimensionality and complexity of the data demand new analytical tools to aid in this task. More precisely, modern neuroscientific data is challenging because: it is high-dimensional, consisting of hundreds of simultaneously recorded neurons alongside rich sensory or behavioral covariates; it is noisy and incomplete, limiting our abilities to draw inferences about single subjects or trials; it is complex, with nonlinear dynamics governing the evolution of neural activity; and it is heterogeneous, with dynamics that differ from one subject to the next.

We construct a class of probabilistic state space models designed to handle these challenging features of neural data, and we develop Bayesian inference algorithms to fit these models at scale. We present these models and algorithms via an illustrative application to population recordings of neural activity in the head ganglia of *C. elegans* collected by Kato et al. [2015] and Nichols et al. [2017].

*C. elegans* is an ideal test-bed for developing new statistical methodologies. The challenges enumerated above appear even in this simple organism, but they are offset by decades of anatomical and behavioral knowledge as well as an expanding experimental toolkit. Advances in optical imaging techniques enable simultaneous recordings of the majority of head ganglia neurons in awake animals responding to sensory stimuli [Schrödel et al., 2013, Prevedel et al., 2014, Kato et al., 2015, Nichols et al., 2017], and these methods have been extended to enable whole brain recording in freely crawling worms [Venkatachalam et al., 2015, Nguyen et al., 2016] where neural activity can be linked to natural behavior [Scholz et al., 2018]. *C. elegans*’ stereotyped nervous system of 302 labeled neurons permits recordings to be aligned across many individuals, ameliorating but not eliminating the difficulty of assembling a canonical model of neural dynamics from many partial recordings. Even in the head ganglia of immobilized worms, neural dynamics are still highly nonlinear and characterized by discrete population states [Kato et al., 2015, Costa et al., 2019]. Understanding how brain dynamics such as these change over the course of development and differ between genetic strains is a fundamental goal of neuroscience. The short life cycle of *C. elegans* and the wealth of genetic tools make it an ideal candidate for studying such questions [Nichols et al., 2017].

We develop a new class of hierarchical recurrent state space models to gain insight into the dynamics of neural activity in the head ganglia of *C. elegans*. We aim to answer four key questions. First, how can we interrogate the *state* of a neural circuit given a collection of relatively short, noisy, and partial recordings collected across multiple individuals? To do so, we build on a long line of work on dimensionality reduction methods for neural data [Yu et al., 2009, Churchland et al., 2012, Cunningham and Yu, 2014, Kobak et al., 2016, Whiteway and Butts, 2016, Wu et al., 2017, Aoi and Pillow, 2018], especially in the case of multiple partial recordings of the same circuit [Turaga et al., 2013, Soudry et al., 2015].

Second, how can we characterize the *dynamics* of these latent states? This has been a central focus of the computational neuroscience community and the broader machine learning, signal processing, and statistics communities for many years. Hidden Markov models (HMMs) [Rabiner, 1989] and linear dynamical systems (LDSs) [Kalman, 1960] are simple, interpretable state space models well-suited to neural data [Smith and Brown, 2003, Jones et al., 2007, Paninski et al., 2010, Macke et al., 2011, Pfau et al., 2013, Chen et al., 2014, Linderman et al., 2016, Gao et al., 2016]. In recent years, these methods have been extended to handle arbitrarily complicated nonlinear dynamics with artificial neural networks [Zhao and Park, 2017, Pandarinath et al., 2018, Hernandez et al., 2018]. We take an intermediate approach, balancing interpretability and flexibility by modeling the dynamics as switching between a handful of simple, linear states [Petreska et al., 2011, Taghia et al., 2018, Wei et al., 2018] according to a switching linear dynamical system (SLDS). We also allow for *recurrent* dependencies between the location of the continuous state and the corresponding linear dynamics [Linderman et al., 2017, Nassar et al., 2019].

Most of the aforementioned methods have focused on multiple recordings of the same population of neurons, where it can be safely assumed that the same dynamics apply to each recording. This is not the case when considering recordings collected from multiple worms, each with their own idiosyncrasies. Our third question is, how can we recover shared structure in the face of such individual and trial-to-trial variability? Hierarchical Bayesian models [Gelman and Hill, 2006] are ideally suited to this challenge.

Hierarchical Bayesian models achieve their flexibility by allowing distinct parameters for each trial, group, subject, etc., and they maintain statistical efficiency by assuming that these low-level parameters share a common prior distribution. For example, hierarchical Bayesian models have been used to model per-neuron tuning curves, assuming each curve is drawn from a shared function class [Behseta et al., 2005, Cronin et al., 2010]; to model trial-to-trial differences in timing [Perez et al., 2013, Duncker and Sahani, 2018], assuming shared response functions after temporal alignment; to “stitch” together non- or partially-overlapping recordings from many individuals, assuming shared underlying dynamics [Soudry et al., 2015, Nonnenmacher et al., 2017, Pandarinath et al., 2018]; and to learn high-dimensional neural responses, assuming shared weights across groups, conditions, and trials [Kobak et al., 2016, Batty et al., 2017, Aoi and Pillow, 2018]. We use hierarchical models to allow each worm its own unique dynamics while still requiring these dynamics to be similar across individuals. In this way, we balance statistical power with model flexibility.

Finally, we ask how the dynamics of neural activity are modulated by external influences and how these effects differ between worms of different age and genetic make-up. For example, oxygen concentration, a salient sensory stimulus for *C. elegans*, is known to modulate neural activity in a manner that depends on developmental stage and genetic strain [Gray et al., 2004, Nichols et al., 2017]. Thus, we introduce input dependencies into the hierarchical model, building on input-output hidden Markov models (IO-HMMs) [Bengio and Frasconi, 1995] to capture the probability of regime switches as a function of external factors.

In the following, we will construct the model, building up to the standard SLDS and then layering in a hierarchical extension, recurrent dependencies, and external inputs. At each stage we show how the increased model capacity captures important features of the data and yields a flexible yet interpretable description of neural population dynamics.

## Results

### Switching linear dynamical system model combines partial recordings from multiple animals to learn shared brain states and dynamics

We construct a model of neural activity in the head ganglia of immobilized *C. elegans* and fit it to partial recordings from multiple worms to learn shared brain states and dynamics. Each neuron in *C. elegans* has a unique cell class identity, such as AVAL or RIVR. The *C. elegans* nervous system is bilaterally symmetric and mostly consists of left/right pairs of neurons. The last letter in the name indicates whether the neuron is the left or right member of the pair. Assigning identities to neurons is a laborious manual task, and only a subset of neurons can be labeled with high confidence. The top row of Fig. 1 shows a two minute window of data for the identified neurons in the five worms studied by Kato et al. [2015]. Across the five worms, 59 neurons were identified in at least one recording. Many other neurons were present in the calcium imaging videos, but they could not be identified with enough confidence to be included in this dataset. Nevertheless, these partial recordings already offer insight into the dynamics of neural activity in the head ganglia.

**Figure 1:**
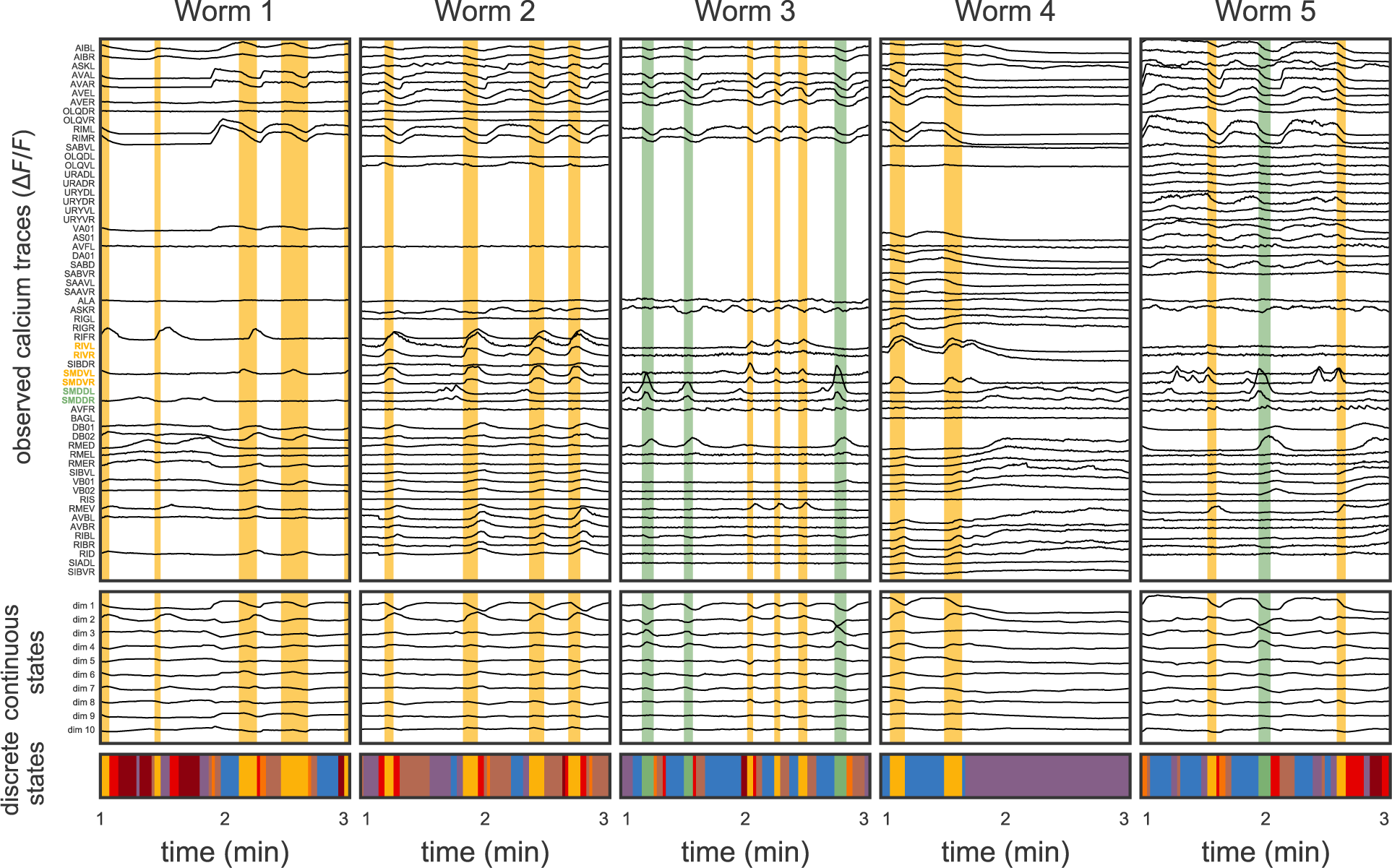
Neural activity reflects discrete and continuous latent states. **Top:** The data consist of partial recordings from many worms. Here we show the calcium fluorescence traces (*ΔF/F*) of the 59 neurons labeled in at least one of the five worms from the [Kato et al., 2015] dataset. For clarity, we show 3 minute segments of the 18 minute long recordings. The neurons are sorted based on similarity (details below). Overlaid in yellow and green are segments of time that our model identifies as instances of two different discrete states. The states correspond to characteristic patterns of population activity, particularly of the six neurons whose labels are colored. These neurons have been implicated in ventral turns (yellow) and dorsal turns (green). **Middle:** We model this population activity as a linear function of a low-dimensional continuous latent state, here a ten dimensional time series found via factor analysis, accounting for the partial observations. **Bottom:** The dynamics of these continuous latent states switch between discrete states, each associated with a different color.

The most noticeable feature in this data is that many neurons have very similar patterns of activity. These similarities suggest that the population’s activity can be summarized with a lower dimensional “continuous state,” as shown in the middle row of Fig. 1. The observed neural activity is modeled as a linear function of the latent continuous state with independent additive Gaussian noise for each neuron, as in factor analysis [Bartholomew et al., 2011]. The continuous latent state evolves over time to reflect the changing activity of the observed neurons and serves as a low dimensional representation of the population activity.

Next, we note that many patterns of neural activity occur repeatedly over time and across worms. For example, the yellow and green bars in Fig. 1 highlight segments of time with stereotyped rises and falls in neural activity and the underlying continuous states. In particular, the yellow segments show rises in the activity of RIVL/R and SMDVL/R, which, in unrestrained worms, are active during ventral turns (reorientations along the ventral body axis) [Kato et al., 2015]. Likewise, green segments correspond to activation of SMDDL/R, which signal dorsal turns [Kato et al., 2015]. In continuous state space, yellow and green segments show characteristic patterns of activity in the first few dimensions. These repeated patterns of activity suggest a simple model of neural activity: the brain switches between discrete latent states, each corresponding to stereotyped changes in the continuous latent states and the observed neural traces. Our fitted model reveals a host of discrete states, as shown in the bottom row of Fig 1. As we will see, these discrete states capture a variety of activity patterns corresponding to different types of behavior.

In this model, each discrete state has a linear dynamics function that determines the temporal evolution of the continuous states. Linear dynamics can capture a variety of activity patterns, like the rises and falls shown above, but they cannot capture more complex patterns of activity. For example, linear dynamics cannot produce multiple fixed points. However, by switching between discrete states with different linear dynamics, the system as a whole can exhibit highly nonlinear dynamics.

This composition of discrete states and linear dynamics is called a switching linear dynamical system (SLDS) [Ackerson and Fu, 1970, Harrison and Stevens, 1976, Chang and Athans, 1978, Hamilton, 1990, Bar-Shalom and Li, 1993, Ghahramani and Hinton, 1996, Murphy, 1998, Fox et al., 2009, Petreska et al., 2011, Linderman et al., 2017, Nassar et al., 2019]. We build upon this model, augmenting it with hierarchical, recurrent, and exogenous input dependencies (Fig. D.1 and Appendix A). We fit these models using a two-stage approach described in Appendix B. Briefly, the SLDS can be seen as the composition of a factor analysis model and an autoregressive hidden Markov model (AR-HMM) [Rabiner, 1989, Hamilton, 1990, Berchtold, 1999] (an HMM with time-dependent autoregressive observations). We first fit the factor analysis model to obtain continuous states, treating the unobserved neurons as missing at random. The continuous states are obtained via *expectation-maximization* (EM) [Dempster et al., 1977], which iterates between computing the expected value of the latent states given the partial observations and finding the most likely linear mapping from continuous states to observed neural activity. Given the continuous states, we infer the discrete latent states and the corresponding model parameters, again using EM. We also develop an end-to-end fitting procedure using black box variational inference [Blei et al., 2017] (Appendix C). Both the two-step and the end-to-end approaches produce an estimate of the underlying continuous and discrete states, as well as the model parameters that govern their dynamics. These reveal interpretable structure in the neural activity of *C. elegans*.

### A low dimensional embedding reveals clusters of functionally similar neurons

The linear mapping from latent states to observed calcium traces reveals clusters of similarly tuned neurons. The mapping is a matrix; rows correspond to neurons and columns correspond to latent dimensions (Fig. 2a). Each row can be seen as an *embedding* that specifies how the corresponding neuron responds to each dimension of the latent state. This matrix reveals groups of neurons with similar embeddings. To capture these groups, we used k-means to cluster the neurons based on the normalized rows of the embedding matrix and found that 10 clusters adequately capture the diversity of the population as measured by the silhouette score (Fig. D.2). Using fewer clusters leads to qualitatively similar groups but omits some of the finer within-group differences; allowing more clusters eventually leads to groups of lateral pairs (like SMDDL and SMDDR in Fig. 2a). These functional clusters were used to order the neurons in Fig. 1.

Fig. 2b shows the cosine similarity of embeddings for each pair of neurons, illustrating the differences and similarities between clusters. We notice two prominent features. First, the block structure of this matrix lends support to the notion of discrete clusters, indicating strong similarity in the activity of neurons within the same cluster in this experimental context. Second, after sorting clusters a block diagonal structure appears, indicating that in addition to the discrete clustering, there is also a continuum along which these clusters vary.

**Figure 2:**
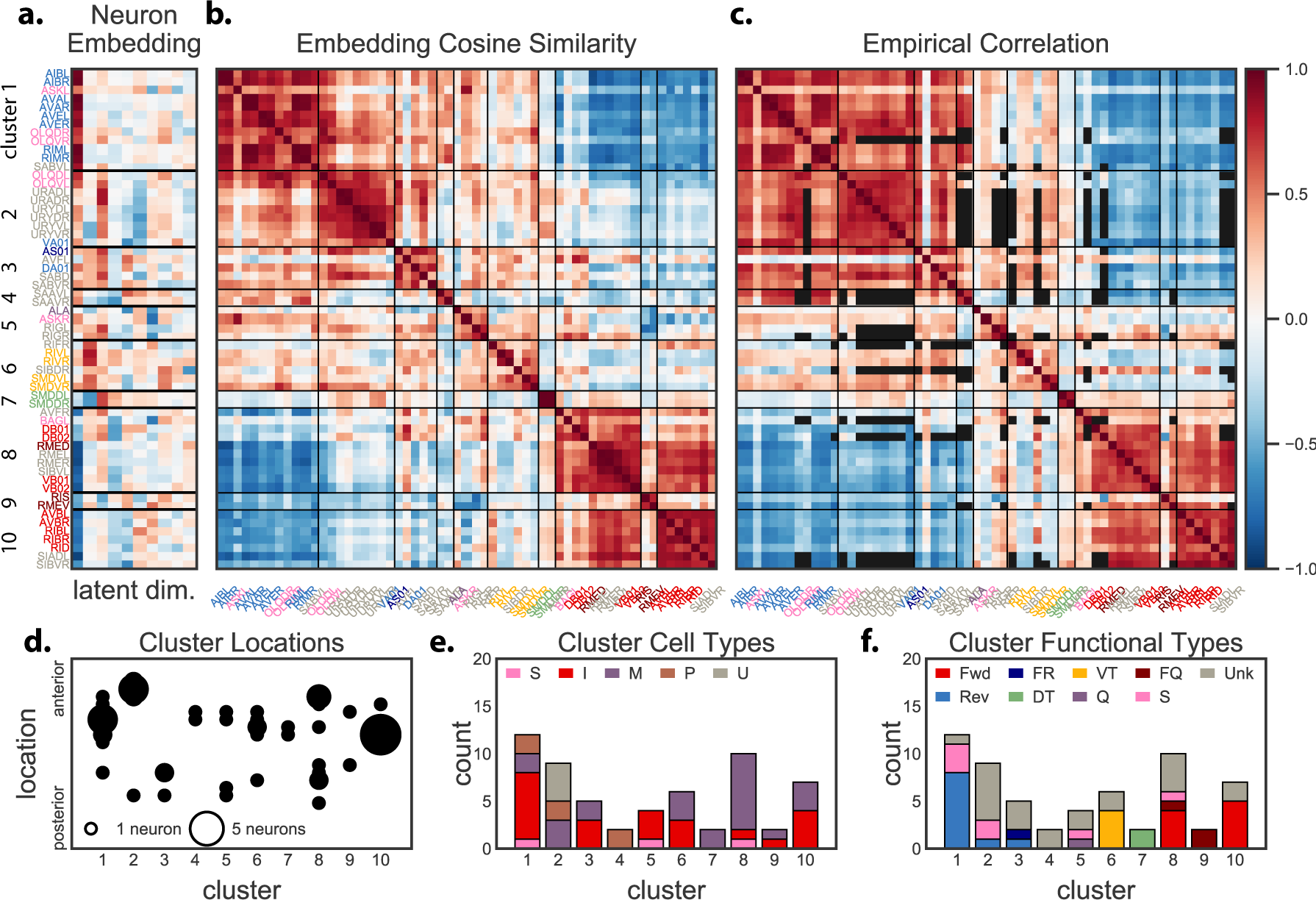
A linear embedding of neurons reveals clusters of functionally similar neurons. **a.** The mapping from continuous latent states to observed calcium traces is linear; we call the matrix implementing this linear mapping the *embedding matrix*. We normalized the rows of this matrix and used k-means to cluster the neurons. Neuron labels are color-coded by their known functions (see panel **f.**). **b.** The cosine similarity of the low dimensional neuron embeddings reflects these clusters and offers a model-based estimate of the correlation between neurons. **c.** By contrast, the empirical correlation is plagued by missing data (black entries) for pairs of neurons that are never simultaneously observed. **d.** The clusters of neurons do not trivially reflect the location of the neurons within the head, as shown by the dispersion in cluster locations. **e.** Nor do the clusters reflect cell type as defined by [Altun et al., 2018], since the clusters are quite heterogeneous (*S: sensory, I: interneuron, M: motor, P: polymodal, U: unknown*). **f.** Instead, the clusters better reflect different functional groups of neurons (*Fwd: forward, Rev: reverse, FR: forward/reverse, DT: dorsal turn, VT: ventral turn, Q: quiescence, FQ: forward/quiescence, S: sensory, Unk: unknown*), as defined in Table E.1.

The cosine similarity of two neurons’ embedding vectors predicts the correlation of their activity. Fig. 2c shows that the predicted correlation matches the empirical correlation matrix. The black cells in the empirical correlation matrix denote missing entries for pairs of neurons that are not simultaneously observed. Accurate estimates of the covariance matrix could facilitate a number of tasks, like predicting the activity of unobserved neurons or aiding in neuron labeling [Linderman et al., 2018].

What can explain the clustering of neurons? We first look at the spatial locations of neurons in each cluster, under the hypothesis that clusters could be trivially explained by an inability to resolve differences in activity between nearby neurons. Fig. 2d shows the locations of all 59 labeled neurons, using the positions from WormAtlas [Altun et al., 2018] along the head-tail axis of the worm’s body. The size of the dot indicates the number of neurons at that location. Some clusters (e.g. cluster 7) are spatially localized, but many clusters contain neurons from multiple head ganglia. This suggests that location alone cannot explain the similarity between tuning vectors.

Next, we considered the types of cells contained in each cluster. Fig. 2e shows the composition of each cluster in terms of sensory (S), inter (I), motor (M), polymodal (P), and unknown (U) neuron types, as specified by the WormAtlas [Altun et al., 2018]. Most clusters are composed of a mix of interneurons and motor neurons, suggesting that cell type alone is also insufficient to explain the clusters.

Specific experiments have shed light on the functional roles of certain neurons. For example, Chalfie et al. [1985] implicated a subset of neurons in forward crawling (e.g. AVBL/R, DB1/2, VB1/2) and reversals (e.g. AVAL/R, AVEL/R, DA1, VA1). Through decades of research, this understanding has been refined and expanded [Gray et al., 2005, Haspel et al., 2010, Kawano et al., 2011, Piggott et al., 2011, Kocabas et al., 2012, Rakowski et al., 2013, Turek et al., 2013, Li et al., 2014, Kato et al., 2015, Lim et al., 2016, Zou et al., 2016, Nichols et al., 2017, e.g.]. We list the known functional roles of some of the head ganglia neurons in Table E.1. Fig. 2f shows the breakdown of clusters by functional role. The clusters are mostly composed of neurons with a common function. The first three clusters contain neurons implicated in reversals; cluster 6 contains ventral turning neurons; cluster 7 consists of two neurons implicated in dorsal turns, SMDDL/R; clusters 8-10 consist primarily of neurons involved in forward crawling. The color scheme in Fig. 2f matches the colors of the neuron labels in panels a-c.

### Discrete behavioral states are automatically identified by their neural dynamics

The dynamics of the continuous latent states suggest that neural activity switches between a handful of discrete states, each with relatively simple linear dynamics. Fig. 3a plots the trajectory of Worm 5’s neural activity in the top two dimensions of the continuous state space, color coded by the discrete state inferred under the SLDS. These are the two dimensions with the highest percentage of explained variance. We see that different states correspond to distinct loops through latent space that recur frequently over the course of the recording. Compare this with Fig. 4b of Kato et al. [2015], where behavioral states were determined based on expert knowledge of how the activity of a core set of neurons relates to known behaviors. For example, rising or high activity of AVA indicates a reversal state, so Kato et al. [2015] and subsequent work by Nichols et al. [2017] used a small manual threshold of the time derivative to detect these rising and high states. In contrast, this probabilistic framework is automatic, combines recordings across worms, and requires no additional preprocessing (e.g., smoothing or derivatives) of the data.

**Figure 3:**
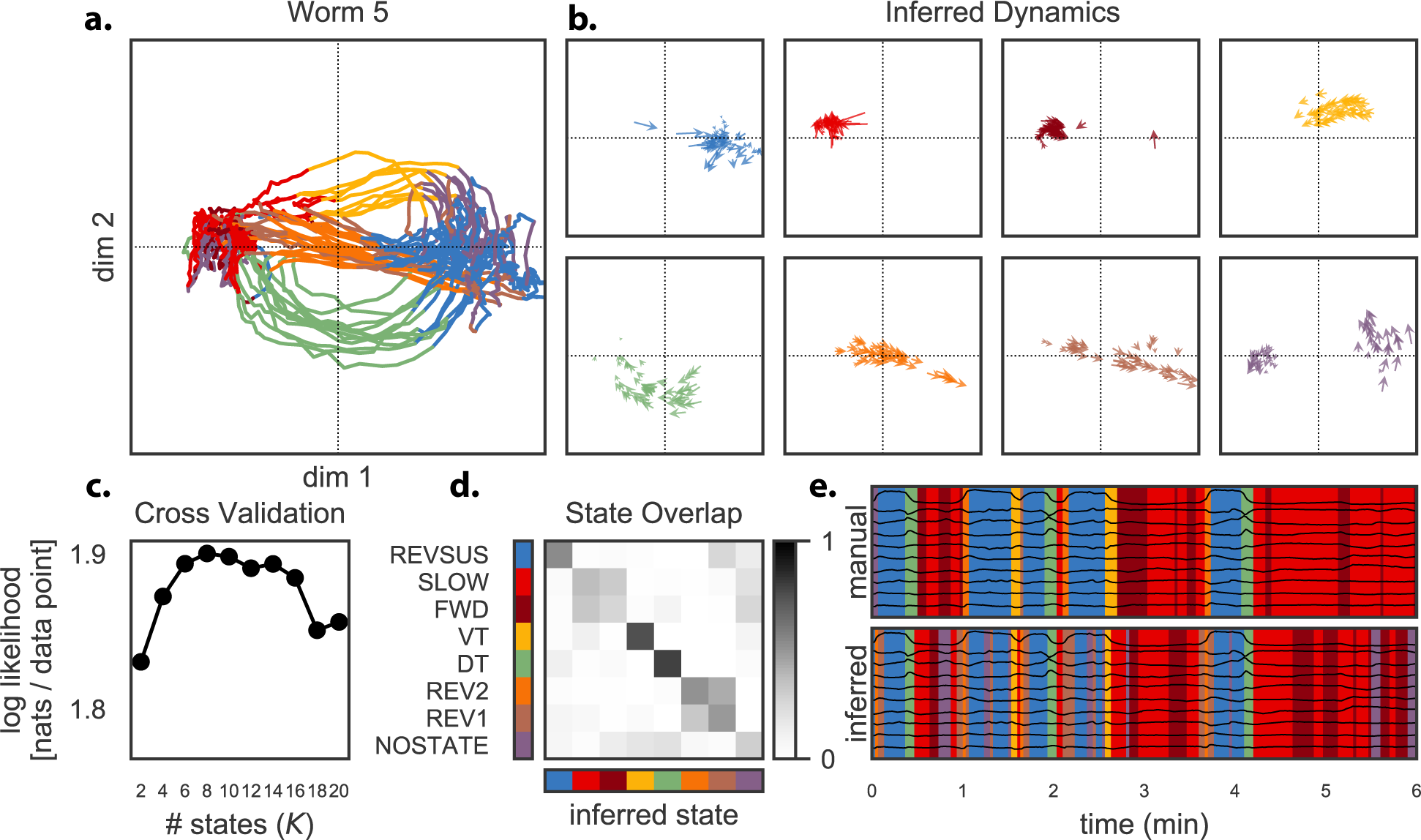
The model segments continuous latent state trajectories into discrete, behaviorally significant states with simple linear dynamics. **a.** The first two continuous latent states, plotted as a trajectory in latent space, trace paths in state space that recur frequently across the recording. The model segments these recurring paths into states with characteristic dynamics, shown here in eight different colors. **b.** Each discrete state (color) is defined by linear dynamics that are used in a particular region of state space. The dynamics can be viewed as vector fields, which show how trajectories move through state space. We show the vector fields at 50 randomly chosen points where the corresponding discrete state was used. Note that the dynamics operate in the full 10-D continuous state space; we show their projection onto the first two dimensions. **c.** The number of states is chosen via cross-validation. The held-out log likelihood plateaus at eight states and then eventually decreases. **d.** The inferred discrete states overlap strongly with the manually labeled states of Kato et al. [2015]. We show the fraction of frames each inferred state was used for a given manual state. Rows sum to one. **e.** This overlap is also shown in the segmentations over time. Here we show the manual and inferred discrete states for the first six minutes of data from worm five.

**Figure 4:**
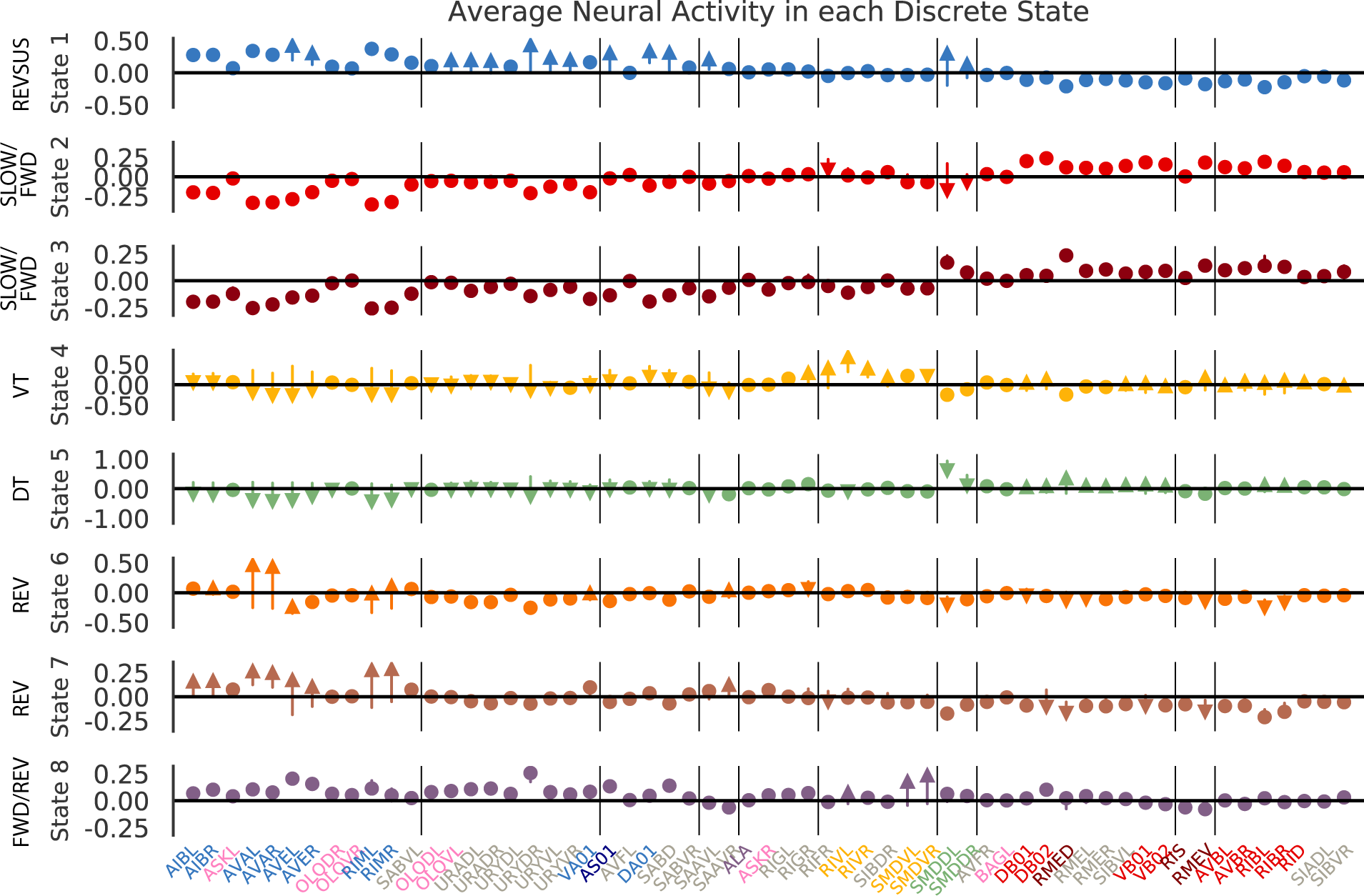
The inferred states activate different subgroups of neurons. Colored symbols show the average change in activity of neurons in each discrete state (i.e. average *ΔF/F* at state exit minus *ΔF/F* at entry). Changes ofless than 0.1 are marked as circles; larger magnitude increases and decreases are marked by arrows. Neurons are sorted and grouped as in Fig. 2. State 1 primarily activates reverse crawling neurons. States 2 and 3 activate forward crawling neurons with very similar average firing rate profiles. States 4 and 5 activate ventral and dorsal turning neurons, respectively. States 6 and 7 are both implicated in reversals, which result in similar increases in the activity of the first cluster of neurons, yet different changes in activation. State 8 activates a mix of forward and reverse crawling neurons.

In the SLDS, each segment is associated with a set of linear dynamics (Fig. 3b). The colored arrows in these vector fields show how the continuous state in one time bin is mapped to a new location in the next time bin according to the corresponding discrete state (for the top two latent dimensions). Arrows are only drawn at those locations where the state was actually employed. Blue, red, and crimson states are all characterized by convergence toward an attractor. These correspond to forward/slowing (red/crimson) and sustained reversal (blue), according to the manual labels below. By contrast, yellow and green trace out trajectories from sustained reverse to forward/slow. These correspond to ventral and dorsal turns, respectively. Orange and brown both correspond to reversals, which link forward crawling to sustained reverse crawling. Purple is the only state that is not spatially localized and limited to a single behavior. Its linear dynamics contribute to the forward/slowing attractor as well as the onset of ventral turns. States like this are rare, but cannot be ruled out entirely.

We use the likelihood of held-out test data to determine the number of discrete states, as shown in Fig. 3c. We see a significant improvement when moving from two to eight states, which matches the number manually chosen by Kato et al. [2015]. Beyond this point, the test likelihood decreases and the additional complexity is not warranted.

Not only does the SLDS find the same number of states, it also finds a correspondence between the manually labeled and inferred states. Fig. 3c quantifies the overlap in inferred states and manually labeled states. Ventral turns (VT) and dorsal turns (DT) are strongly overlapping in manual and inferred states. Some manual states are combined. For example, reversals (REV1 and REV2) are combined, as are forward (FWD) and slowing of forward crawling (SLOW. Fig. 3d shows manual and inferred segmentations over time for a six minute section of data in Worm 5. For visualization, we added a penalty on state transitions (specifically, we added ten times the identity matrix to the log transition matrix) before computing the inferred segmentation in order to limit rapid switching. The manual and inferred states show clear similarities but also some differences. To better understand their differences, we overlaid the continuous states. We see that inferred transitions often correspond to fluctuations in continuous states that are overlooked by the manual segmentation rules. Overall, the correspondence between manual and inferred states shows that the SLDS is a useful tool for automatically segmenting brain data into behaviorally meaningful states.

### Discrete states activate different clusters of neurons

Fig. 4 shows the average activity of each neuron in each discrete state. We organized neurons into clusters, as in Fig. 2, using the same color scheme for neurons based on their known functional roles. We see that the discrete states tend to correspond to activation of certain clusters of neurons; for example, State 1 (blue) has high activity in the first cluster of neurons, which are implicated in reverse crawling. States 2 and 3 (red and crimson) have high activity in forward crawling neurons (red and crimson). State 4 (yellow) coincides with the manually labeled “ventral turn” state. Its dynamics produce a pronounced rise and fall in the activity of RIVL/R and SMDVL/R neurons. The RIV pair are polymodal interneurons and motor neurons that specify the ventral bias of turns, and the SMDV are motor neurons that initiate ventral turns. Likewise, State 5 (green) corresponds to positive activation of SMDDL/R, which start rising at the end of prior sustained reversal and then fall after transition to DT state.

Other states are best understood by the corresponding change in activity they produce. For example, States 6 and 7, which overlap with the reversals identified by Kato et al. [2015], give rise to a large increase in the activity of reverse crawling neurons. In contrast, States 4 and 5, which correspond to turns, show large decreases in the activity of reverse crawling neurons and increasing activity in forward crawling neurons. State 1 shows a small increase in the second cluster of neurons (URAD/V, URYD/V, etc.), whose functional roles could not be confidently asserted. This change in activation implies that these neurons show increasing activity during reversals.

### Hierarchical models capture shared structure while allowing for individual differences between worms

A major challenge in analyzing modern neural datasets is that recordings are often made over the course of many sessions, trials, or animals. We want to combine these disparate recordings to learn a shared model of the system under study, but in order to do so we must account for discrepancies and differences between recordings. This challenge is particularly evident in our study of *C. elegans*, where we want to combine recordings across multiple worms. Each worm differs slightly from the next in ways that are challenging to control experimentally (e.g., the level of GCaMP expression in individual neurons), but they may also differ slightly in terms of their underlying neural dynamics. We expand the model to handle these worm-to-worm differences while still capturing properties that are shared by all worms.

We allow for individual and/or trial-to-trial variability by introducing a hierarchical prior on the parameters of the continuous and discrete dynamics models. Specifically, we model each worm’s dynamics parameters as conditionally independent random variables that share a global mean (Appendix A.2). The global mean captures the discrete states and dynamics of a “canonical” worm, and the per-worm parameters capture the unique properties of individual worms differ. The hierarchical model regularizes the per-worm parameter estimates, biasing them toward the global mean. This regularization leads to more stable estimates, especially given noisy and limited data.

One consequence of the hierarchical model is that state segmentations are more consistent across worms. We see this in Fig. 5a, where the color-coding (i.e. the segmentation) of the continuous trajectories is largely conserved across the five worms, even though the particular shape of the trajectories differs slightly from one to the next. Without the hierarchical model, slight differences in trajectories are segmented into different discrete states, even though by eye they seem to play the same functional role. This can be seen in Fig. 5b. For example, the yellow state (ventral turn) is split into multiple states (yellow, pink, and gray) by the non-hierarchical model.

**Figure 5:**
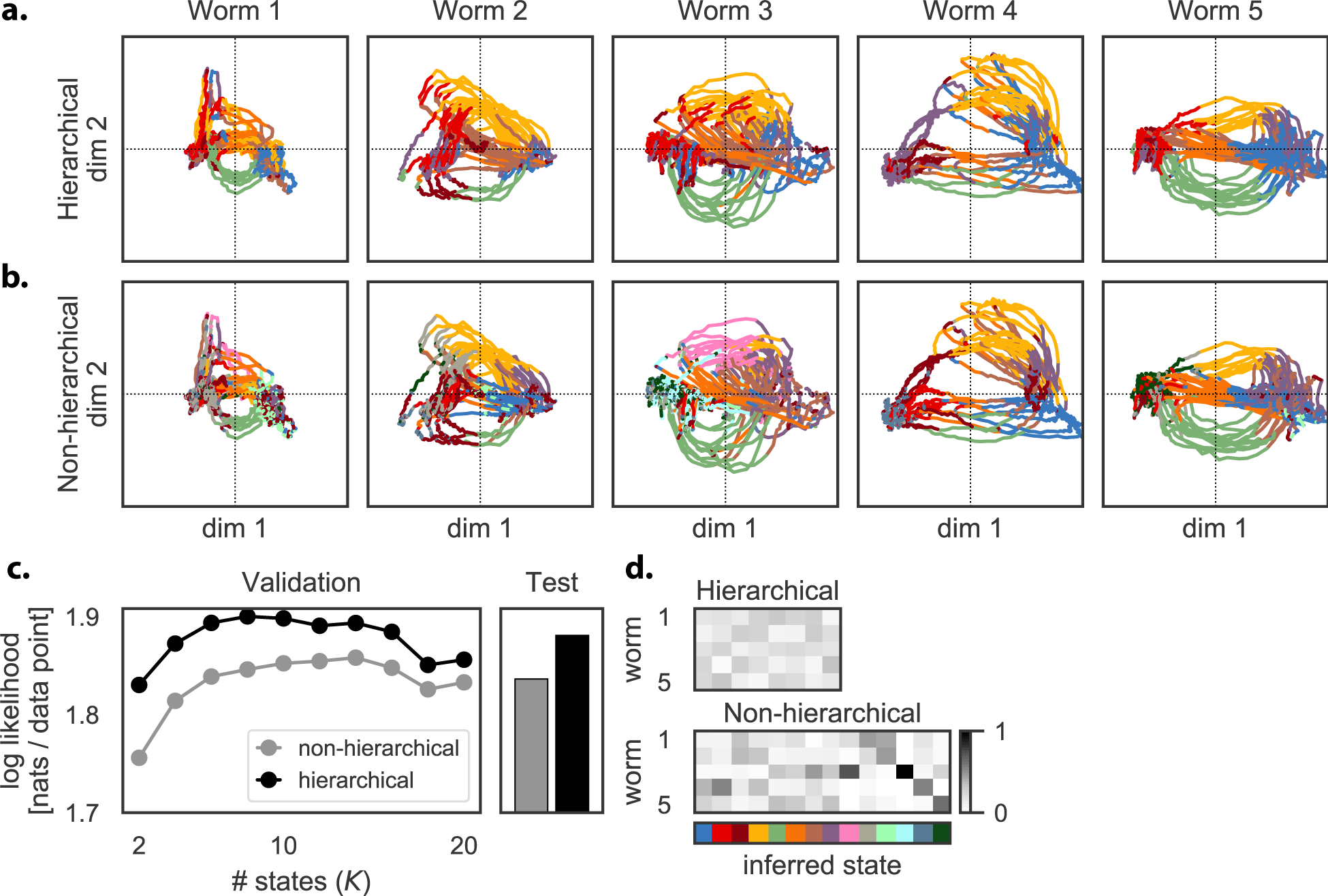
The hierarchical model captures canonical dynamics while allowing for worm-to-worm variability. **a.** Individual latent trajectories differ in shape from worm to worm. For example, in Worm 4, the ventral turn loop (yellow) is exaggerated and the dorsal turn loop (green) is compressed relative to other worms. Still, the overall discrete state sequences are preserved: each worm cycles between forward and reverse states linked by turn and reversal states. The hierarchical model captures this shared discrete structure despite differences in the trajectories from one worm to the next. **b.** The standard model requires state dynamics and hence trajectories to be the same in all worms. Thus, when given the same continuous trajectories as in **a.**, the standard model creates more discrete states in order to handle the idiosyncrasies of each worm. **c.** Hence, the standard model requires 14 discrete states rather than 8, and it still achieves a smaller likelihood on held-out data. **d.** Worm-specific states are evident in the relative usage of states across worms in the hierarchical and standard model. (Columns are normalized.)

Over-segmentation is also evident in the number of discrete states chosen by cross validation, as shown in Fig. 5c. The best hierarchical model has 8 states whereas the best non-hierarchical model has 14; however, even with more states, the non-hierarchical model is less accurate on both validation and test data. Finally, we see in Fig. 5d that the states of the hierarchical model are used much more uniformly across worms than are those of the non-hierarchical model. In this dataset, the five isogenic worms are expected to exhibit the same set of canonical behaviors and hence the same gross patterns of neural activity. Thus, we prefer the simpler description offered by the hierarchical model not only because it achieves higher held-out likelihood, but also because it groups intuitively equivalent discrete states together across worms, even though their dynamics differ slightly.

### Recurrent models learn spatially localized states and transition probabilities

To accurately model the transitions between states, the changes must occur at specific boundaries in continuous latent space. In the hierarchical recurrent SLDS, two factors determine the transition probabilities: the Markov transition weights, which capture the probability of switching from one discrete state to another, and the recurrent weights, which modulate transition probabilities as a function of location in continuous state space. The Markov and recurrent weights are combined via a generalized linear model with a “softmax” function to output a distribution over the next discrete states (Appendix A), as in an IO-HMM [Bengio and Frasconi, 1995] or a recurrent SLDS [Linderman et al., 2017]. The combination of past discrete and continuous states allows the model to capture transition probabilities that change as a function of continuous states.

The nonlinear function that maps states to transition probabilities makes the raw weights less interpretable. Instead, we examine statistics of the transitions. The empirical transition probabilities, obtained by counting transitions under the automatic state segmentations shown in Figures 1 and 3 and neglecting self-transitions, show the common transition patterns first observed by Kato et al. [2015]. Sustained reverse crawling states (blue and purple) transition to forward crawling (red/crimson) via dorsal and ventral turns (green and yellow). The return sustained reversal is made via reversal states (orange and brown).

The recurrent weights modulate the transition probability based on the location in continuous state space. The shading in Fig. 6b indicates the probability of staying in each discrete state for each location in the first two continuous state dimensions. The recurrent weights effectively specify when to leave a discrete state and enter a new one. This is evident from the overlaid trajectories, which are randomly sampled from all segments of the discrete state: many trajectories (e.g. yellow, green, orange, and brown) end near a boundary dictated by the recurrent weights. By combining recurrent weights and Markov transition weights, we obtain a model that approximates nonlinear dynamical systems by probabilistically switching between linear dynamical states in a location-dependent manner. In many locations, the probability of staying in the same state is nearly one, indicating that the discrete transitions are nearly deterministic. As we will see next, recurrent transition probabilities are also key to building more accurate generative models of neural activity.

**Figure 6:**
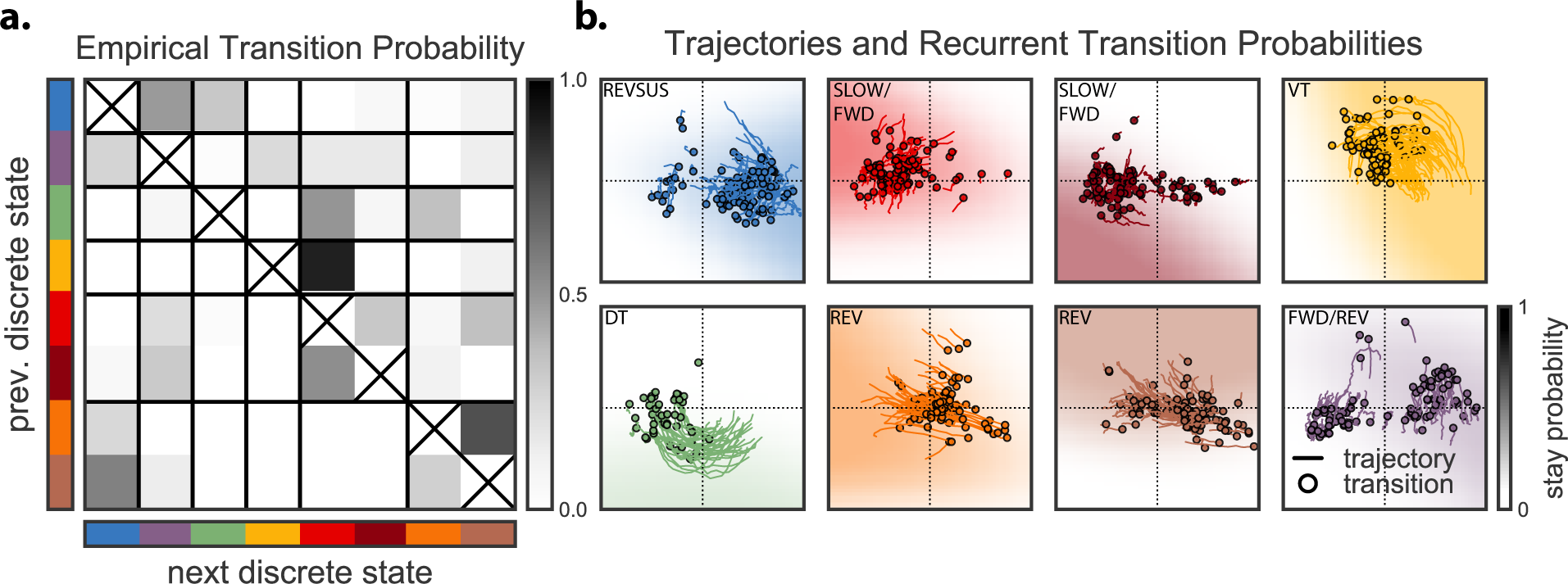
Recurrent models use both the discrete and continuous latent states to determine a probability distribution over the next discrete state. **a.** The normalized transition counts (ignoring self transitions) under the inferred discrete state sequence show a pattern of transition from sustained reversals (blue) to dorsal and ventral turns (green and yellow) to forward crawling (red and crimson) and back via reversals (orange and brown). **b.** The transition probabilities are also determined by “recurrent” weights, which modulate transition probability as a function of location in continuous state space. Here we show the probability of staying in a given state (background shading) as a function of location in the first two dimensions of state space. In many locations, the discrete state transitions are nearly deterministic. For reference, we show 100 illustrative trajectories from entrance to exit transition (marked by a black circle) of the corresponding state. The transition locations for some states, like yellow, green, and orange, are largely determined by the recurrent weights, since those locations often occur in regions where the stay probability has diminished.

### Hierarchical recurrent SLDS are effective generative models of neural population data

We have shown that the hierarchical and recurrent SLDS decomposes neural activity into discrete states with simple linear dynamics, that these discrete states correspond to different elements of the worm’s behavioral repertoire, and that the transition probabilities are informed by both the preceding discrete state and the location in continuous state space. But have we constructed a good model of neural activity? To answer this question, we compare the real neural activity to simulated neural activity generated by the probabilistic model.

First, note that the segmentation tasks considered so far only use the posterior distribution over discrete states under the learned linear dynamics and transition probabilities. It is possible to obtain segmentations that align with known behavioral states without a good generative model. In fact, simple mixture models, which do not model temporal dynamics at all, achieve similarly high overlap with the manually labeled states (Fig. D.3). The reason simple models may suffice for segmentation is that, here, the posterior distribution is dominated by the likelihood, i.e., the linear dynamics model. Ignoring the discrete transition probabilities may not change the posterior distribution substantially, as long as the likelihood is accurate.

On the other hand, to generate realistic synthetic neural traces, learning an accurate discrete state transition model is just as important as learning accurate linear dynamics. Transitioning to a new discrete state at the wrong location in continuous state space will yield unrealistic, and often divergent, trajectories. This is evident in Fig. 7. We show a six minute segment of real neural activity from Worm 5 for reference (Fig. 7a). Alongside, we show simulations from the standard model (Fig. 7b) and from the hierarchical, recurrent model (Fig. 7c). Below we show the continuous and discrete states (inferred for the real data; generated for the simulations). The differences are readily apparent in the low dimensional state space. Here we show the inferred continuous trajectories for each of the five worms (Fig. 7d) alongside simulations from the standard model (Fig. 7e) and the hierarchical recurrent model (Fig. 7f). The standard simulations fail because they switch between discrete dynamical states in a way that is independent of the continuous location. Over the course of an 18 minute simulation, they generate trajectories that switch erratically between discrete states and often diverge in continuous state space. By contrast, the recurrent model transitions at appropriate locations and produces trajectories that are visibly more similar to the real data.

**Figure 7:**
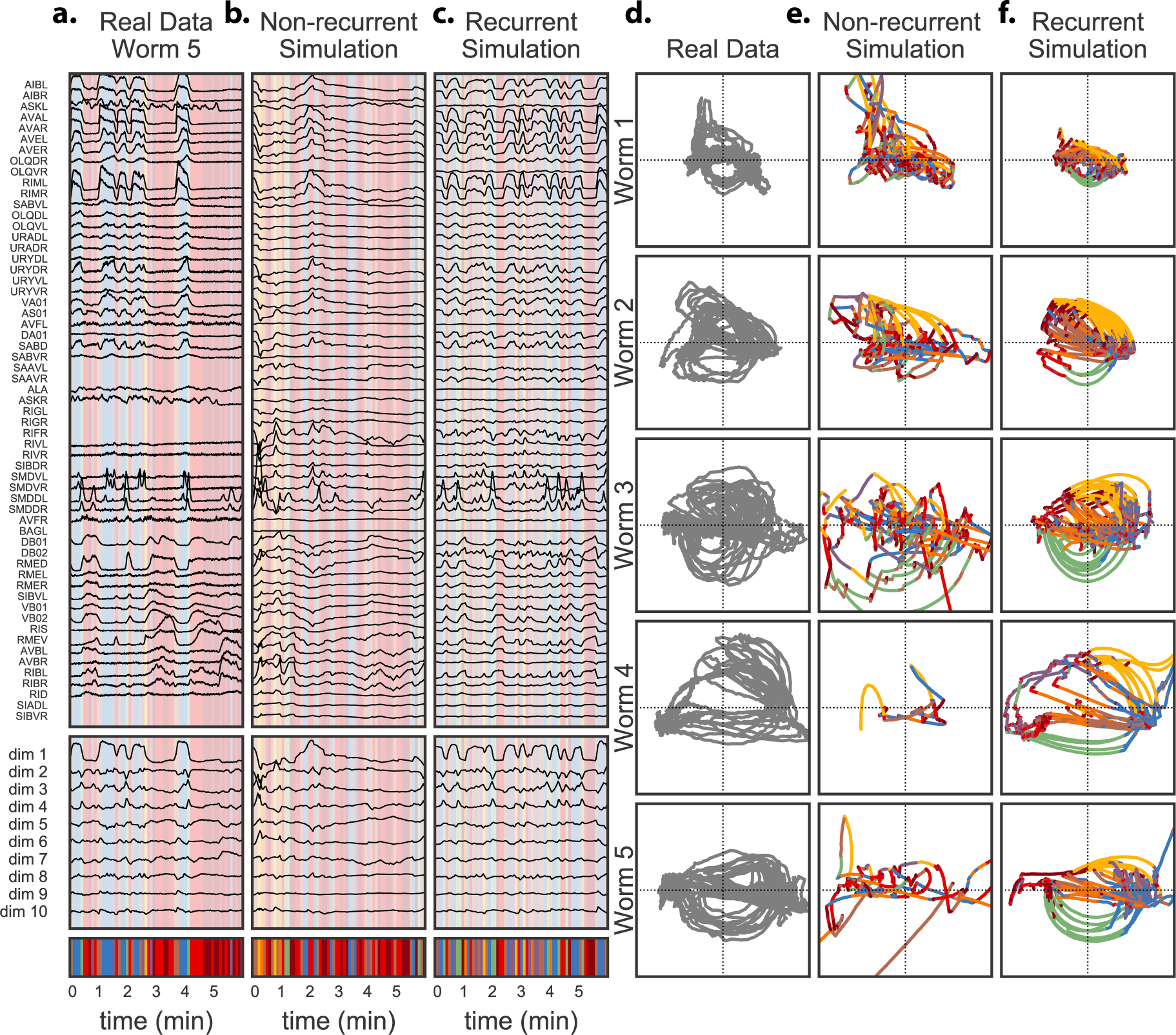
The hierarchical recurrent model generates more realistic neural trajectories. **a.** Real neural trajectories from one worm, along with the inferred continuous and discrete latent states. **b.** Simulations from the non-recurrent model are highly unrealistic and unstable because the discrete transitions are independent of the location in continuous state space. **c.** In contrast, simulations from the recurrent model, which have both discrete and continuous dependencies, recapitulate the salient features of the real neural activity. **d.** For reference, we show the continuous latent states of all five worms. **e.** The 18 minute non-recurrent simulations, viewed as trajectories in latent state space, switch at random times and produce unrealistic and often unstable dynamics. **f.** The 18 minute hierarchical recurrent model is able to capture realistic trajectories and transition times and locations, and thereby simulate realistic activity patterns. Eventually it also becomes unstable, but still produces more realistic trajectories than its non-recurrent counterpart.

### Oxygen modulation of transition probabilities depends on genetic strain and developmental stage

The preceding results demonstrated the efficacy of the hierarchical, recurrent SLDS model for learning interpretable dynamics of neural activity from the noisy, partial recordings of Kato et al. [2015]. We now turn to the dataset studied in Nichols et al. [2017]. There are two genetic strains of worms in this dataset: the standard laboratory strain, N2, as well as N2-derived worms with a loss-of-function mutation in the neuropeptide receptor gene (*ad609*), henceforth *npr-1* worms. As *C. elegans* prepare to exit any of their four larval stages they enter an approximately 2 hour long state called lethargus. Here the animals display behavioral quiescence which has been defined as a sleep state [Raizen et al., 2008]. Nichols et al. [2017] studied larval stage 4 animals before this lethargus state (henceforth, prelethargus) and animals in lethargus. In total, the dataset contains 44 worms: 11 N2 prelethargus, 12 N2 lethargus, 10 *npr-1* prelethargus, and 11 *npr-1* lethargus worms. In each worm we have between 22 and 42 identified neurons, with a median of 35 identified neurons per worm. Moreover, Nichols et al. [2017] modulate the oxygen concentration over the course of each recording to elicit different neural responses. We show how hierarchical recurrent state space models can be extended to study the strain- and stage-dependent effects of these manipulations.

Fig. 8a shows the five dimensions of the continuous states that capture the most variability in the neural activity, averaged over the 10-12 worms in each group. Fig. D.4 shows the raw continuous traces from which these averages were computed. We see that prelethargus worms exhibit sustained neural activity throughout the recording session, regardless of O_2_ concentration; lethargus worms, by contrast, show less neural activity, especially at 10% oxygen concentration. As Nichols et al. [2017] showed, the neural activity of N2 and *npr-1* lethargus worms differs when oxygen concentration is increased to 21%: higher oxygen concentration elicits sustained neural activity for *npr-1* worms, but only a transient (1-2 minute) response in N2 lethargus worms. Fig. D.5 shows examples of raw neural traces (ΔF/F) for each group.

**Figure 8:**
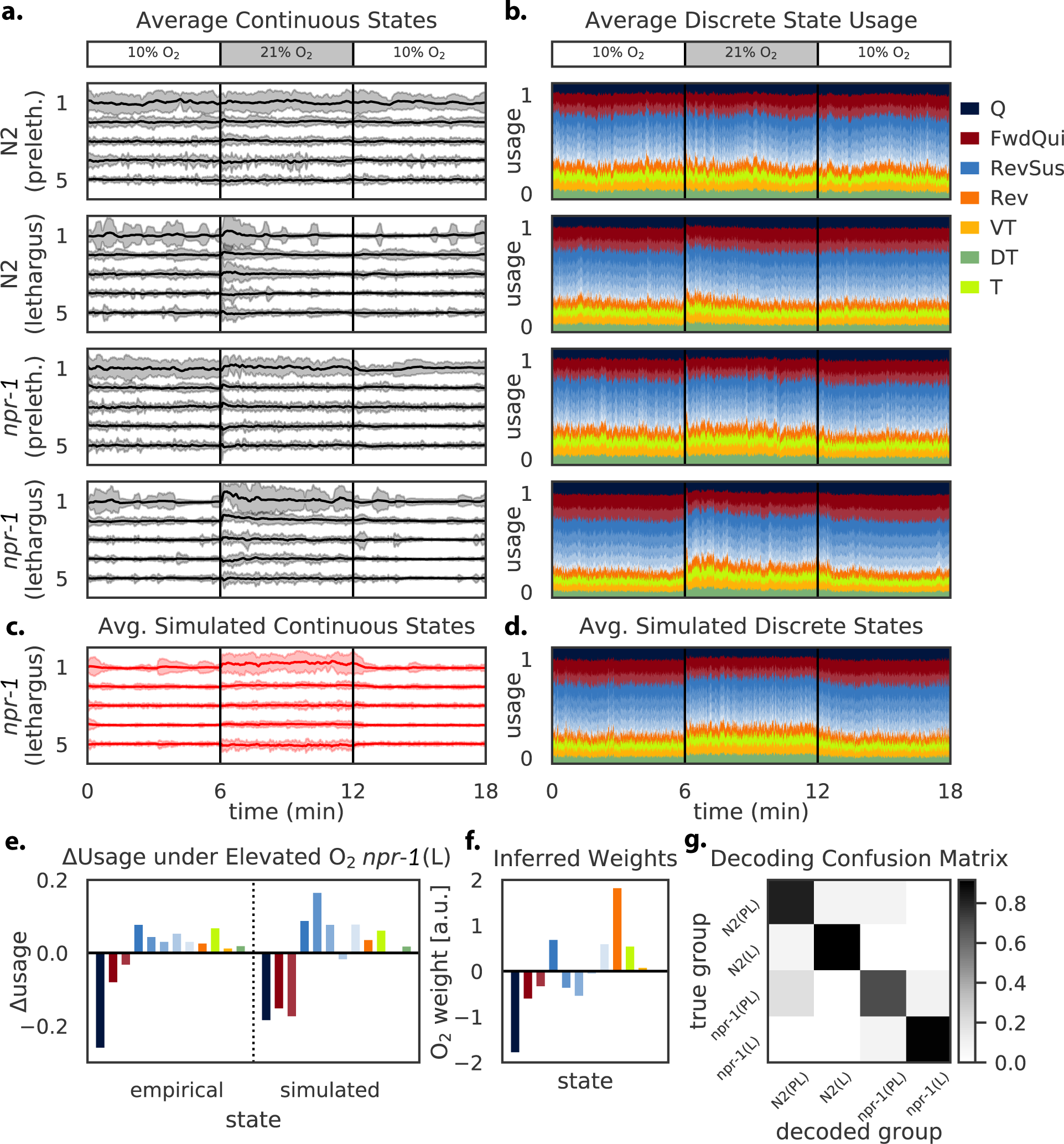
Exogenous inputs such as oxygen concentration modulate transition probabilities differently across genetic strains and developmental stages. **a.** Nichols et al. [2017] measured neural activity in 44 worms spanning two genetic variants (N2 and *npr-1*) and two developmental stages (lethargus and prelethargus). Each worm is subject to time-varying O_2_ concentration (top). **a.** Average activity in the top 5 (of 10) continuous state dimensions for each group (*n* =10-12 worms per group). Shading denotes three standard deviations. As Nichols et al. [2017] observed, prelethargus worms are relatively insensitive to O_2_ concentration change. Lethargus worms respond to elevated O_2_ in a strain-specific manner: N2 worms exhibit a transient increase in activity (1-2 minute increase in mean and variance), whereas *npr-1* worms show a sustained increase throughout the 6 minutes of elevated O_2_. **b.** The hierarchical recurrent SLDS captures these changes in activity with a shift in discrete state usage, shown here as a “mountain” plot indicating the average state probability over time for each group (with 2s Gaussian filter). Colors match preceding plots with addition of *quiescent* (Q) and a general *turn* (T) states. Multiple discrete states appear to map to forward/quiescent (FwdQui) and sustained reversal (RevSus); we distinguish these states with shades of the same color. **c-d.** We capture this oxygen dependence by adding exogenous inputs to the model with separate input weights for each group. Simulations from the fitted model recapitulate the average continuous discrete and continuous state usage across groups, shown here for only *npr-1* lethargus worms. **e.** The *npr-1* lethargus worms exhibit an increased usage of reversals (orange/brown), sustained reversal (blue), and turning (chartreuse, yellow, green) over the period of elevated O_2_ and a decrease in forward crawling and quiescence. **f.** The model captures this change in usage primarily through a decrease in quiescence and an increase in the probability of reversals, which are then followed by sustained reversal and turns under the transition model (c.f. Fig 6). **g.** Finally, using the inferred input dependencies, we can decode which group a worm belongs to on the basis of its observed neural activity in response to fluctuations in oxygen concentration.

The hierarchical recurrent SLDS explains this increased variability in continuous states via an oxygen-dependent shift in discrete state usage. Fig. 8b shows the average inferred discrete state usage over time for each group; this is obtained by taking the most likely state sequence for each worm, computing the empirical probability of each discrete state within each time bin for each group, and smoothing with a 2 second Gaussian filter. Colors match those in preceding figures with the addition of a “quiescent” state (Q), a forward/quiescent state (FwdQui), and a general “turn” state (T) when direction is ambiguous. As before, the discrete state labels are determined by assessing the average activity and change in activity of each neuron in each discrete state (Fig. D.6). With this larger and more diverse dataset, cross-validation yields 12 discrete states. Some of these states have similar patterns of neural activation (Fig. D.6), so we label them as separate instances of the same behavioral state and distinguish them with different shades of the same color. Moreover, the distinction between forward and quiescent states is subtle—it is determined by the activation of RMED, RMEV, and RIS, which are active during both forward and quiescent behavior [Turek et al., 2013, Nichols et al., 2017]. The quiescent state corresponds to activity in these three neurons but not in the general forward crawling neurons, whereas the forward/quiescent states show activity in these three neurons as well as the forward crawling neurons (Fig. D.6).

Prelethargus worms show limited change in discrete state usage in response to elevated oxygen concentration. Lethargus worms of both strains show an initial decrease in quiescent state usage upon increase in oxygen concentration, but only the *npr-1* variants show a prolonged decrease in quiescent states throughout the 6 minutes of elevated oxygen concentration.

To capture these exogenous input dependencies, we construct an input-driven hierarchical recurrent model in which the instantaneous oxygen concentration modulates the transition probabilities. Fig. D.1d shows its graphical model, where the input is the absolute oxygen concentration. The input combines linearly with the Markov and recurrent weights to determine the log probability of discrete state transitions. Following the hierarchical models developed above, we give each group its own input weights, and each individual worm its own Markov weights, recurrent weights, and linear dynamics. As before, the hierarchical model penalizes differences between individual parameters and the global mean. Weakly tying the parameters allows us to learn per-group and per-individual parameters while sharing statistical strength across the entire population of worms. See Appendix A.2 for complete details.

The fitted model recapitulates many patterns of continuous and discrete state dynamics. Fig. 8c shows the continuous state usage for simulated *npr-1* lethargus worms averaged over *n* = 10 samples from the generative model. As in the real data above, we see an elevated mean and variance in activity along the first continuous state dimension at 21% O_2_. The patterns of *npr-1* lethargus discrete state usage are recapitulated in the simulated data shown in Fig. 8d. Fig. 8e summarizes the change in state usage at 21% O_2_ by taking the difference in average state usage in minutes 6-12 versus 0-6 and 12-18. Again, the increase in reversals, sustained reversals, and turns at the expense of quiescent and forward crawling patterns of neural activity is visible in both the real and simulated data. These broad changes in state usage are achieved with a sparse change in the transition probabilities, as shown by the inferred input weights in Fig. 8f. In the fitted model, increased O_2_ concentration leads to an increase in the probability of reversals (orange bar), but a relatively minor change in the probability of other states. Since reversals lead to sustained reversal and eventually to turns (c.f. Fig. 6), the increase in reversals leads to the broader changes in state usage seen in Fig. 8e.

Finally, having developed and fit a model of how neural dynamics in the head ganglia respond to oxygen concentration across multiple genetic strains and developmental stages, we can use the model to *decode* the strain and stage of the worm from observed neural activity. We evaluated the likelihood of each worm’s neural activity, holding its per-worm parameters fixed and swapping the input-weights from each of the four groups. The decoded group is the one which assigns highest log probability to the neural data. As shown in Fig. 8g, this decoding procedures identifies the correct group with high probability, suggesting that hierarchical, recurrent, and input-driven state-space models are well-suited to not only generating realistic neural data, but also for quantifying the differences in neural population activity across genetic strains and developmental stages.

## Discussion

A fundamental goal of systems neuroscience is to understand how sensory inputs modulate neural activity and how this neural activity ultimately drives natural behavior. *C. elegans* is an ideal model organism in which to pursue this goal: its nervous system is stereotyped across individuals and well-mapped, optical tools enable large-scale measurement of neural activity during unconstrained movement, and its behavior is nontrivial but not overly complex. In order to capitalize on this potential, however, we need new computational and statistical tools. The model we developed here can identify salient dimensions of neural activity, quantify how the brain activity dynamics differ across individuals, and capture the relationship between sensory inputs, neural dynamics, and behavioral outputs.

Discrete decompositions are an attractive method for simplifying nonlinear brain dynamics. Recently, SLDS models have been applied in a variety of neural modeling tasks, particularly for segmenting recordings of brain activity into behaviorally relevant or task-relevant epochs [Petreska et al., 2011, Taghia et al., 2018]. Similar techniques have been developed specifically for modeling *C. elegans*, and have also studied the data from Kato et al. [2015] and Nichols et al. [2017]. Costa et al. [2019] develop an adaptive, locally-linear model of neural activity in which dynamics can change at any point in time, and they fit the model using a change-point detection algorithm based on a likelihood ratio test. We share the same goal, but instead take a Bayesian hierarchical modeling approach, which allows us to compare various extensions to the probabilistic model and appeal to a host of Bayesian inference algorithms. Brennan and Proekt [2017] take a slightly different approach, first characterizing the low-dimensional manifold of neural activity, then breaking it down into “loops.” To model dynamics on the manifold, they keep track of the current loop and the “phase” along this loop. This approach is analogous to keeping track of a discrete state and the time since transitioning into it, as in hidden semi-Markov models [Murphy, 2002, Yu, 2010, Johnson and Willsky, 2013]. An interesting avenue for future work is to consider how semi-Markov and recurrent transition models can be combined to better model state transitions and durations.

We applied the hierarchical, recurrent, and input-driven model to immobilized worms and showed that we can automatically recapitulate salient patterns of neural activity and their relationship to environmental stimuli, genetic strain, developmental stage, and known behavioral primitives. There are some clear next steps. First, these methods could be extended to understand how neural dynamics are shaped by more complex multi-sensory stimuli and broader sets of genetic and neural perturbations. Likewise, these models are ready to be deployed on measurements of neural activity in unconstrained worms. Immobilized worms likely express a limited repertoire of behavioral commands. Therefore, a fundamental question for future study is whether the full variety of worm behaviors requires a larger repertoire of underlying latent discrete states or greater control over local dynamics. Addressing these problems could lead to new insights into how behavior might be organized hierarchically and across time-scales. One of the limitations to measuring neural activity in freely crawling worms, however, is the difficulty of tracking neurons through time and space and resolving their identities. Soon this task will be improved and accelerated with the advent of multi-color imaging techniques that assign a unique color to each neuron [Yemini, 2018], and machine learning methods that combine color and location information to find an optimal assignment of names to neurons [Linderman et al., 2018, Mena et al., 2018].

Recurrent SLDSs offer a statistical description of neural time series, but they may also suggest underlying principles of neural computation. For example, the fundamental assumption of this model is that neural dynamics arise from the combination of simple, linear dynamical regimes along with a set of state-dependent rules for transitioning between them. Switching between linear dynamical regimes gives rise to highly nonlinear dynamics. The recurrent SLDS assumptions stand in contrast to classical recurrent neural network models, in which the dynamics are inherently nonlinear [Dayan and Abbott, 2001]. There are several possible advantages to a modular decomposition, separating transition rules from low-level dynamics. Modular systems may be easier to control, allowing exogenous inputs to vary the probability of a discrete set of states (as in Fig. 8f) rather than the continuous space of neural activity. This potentially could enable animals to generate robust behavioral patterns in the presence of complex multi-sensory inputs, which in this view primarily control transitioning between otherwise robust dynamical regimes. Future work, employing more complex stimulus-paradigms can address these questions. Moreover, the influence of location within state on transition probabilities provides a means to predict variability and context dependence in sensory responses [Gordus et al., 2015, Liu et al., 2018].

Systems that are simpler to control may also be easier to learn, adapt, or evolve, since the dynamics can be held fixed while the transition probabilities are varied. Finally, we note that recurrent networks with rectified linear units also lead to piecewise linear dynamics models, albeit with far more linear regimes. The fact that the recurrent AR-HMM recapitulates neural activity and finds a small number of discrete states that correspond to natural elements of worm behavior suggests that this modular decomposition provides a useful statistical description of *C. elegans* neural dynamics. However, understanding how neural circuitry in *C. elegans* implements these dynamics remains an open question.

State space models address break down complex neural data into a sequence of discrete and continuous states. In *C. elegans* we have a unique opportunity to study the mechanistic bases of these states by relating them to the synaptic connectivity between pairs of neurons. Our ability to do so with current data is limited by the fact that we only observe a subset of all 302 neurons, but we expect that improved recording technologies will soon overcome this challenge and allow truly whole brain recordings with adequate temporal and spatial resolution. The methods developed here will then provide an intermediate target for relating low-level implementation to high-level behavioral outputs, mediated by discrete and continuous states of neural activity.

## Acknowledgments

S.L. is supported by a Simons Collaboration on the Global Brain postdoctoral fellowship (SCGB-418011). D.B. is supported by ONR (N00014-17-1-2131, N00014-15-1-2209), NIH (1U01MH115727-01), and DARPA (SD2 FA8750-18-C-0130). The research leading to these results has received funding from the European Community’s Seventh Framework Programme (FP7/2007-2013)/ERC grant agreement 281869, and the Research Institute of Molecular Pathology (IMP) (A.L.A.N, M.Z.). M.Z is supported by the Simons Foundation (#543069) and the International Research Scholar Program by the Wellcome Trust and Howard Hughes Medical Institute (#208565/Z/17/Z). The IMP is funded by Boehringer Ingelheim. L.P. is supported by the Simons Collaboration on the Global Brain and a NSF Neuronex Award (#1707398).

## A Hierarchical recurrent state space model

We model the neural activity with a switching linear dynamical system (SLDS) with three extensions: (i) a *recurrent* model to capture how continuous latent states influence discrete state transition probabilities; (ii) a *hierarchical* model to share parameters across worms while also allowing for worm-to-worm variability; and (iii) an *input* model for capturing the effects of exogenous factors, like oxygen concentration, on discrete state dynamics. The graphical models corresponding to the standard SLDS and its extensions are shown in Fig. D.1, and Table A.1 summarizes the notation used in this section. We will introduce the SLDS first and then present each of these extensions in turn.

### A.1 The standard SLDS

Switching linear dynamical system models (SLDS) break down complex, nonlinear time series data into sequences of simpler linear dynamical modes. By fitting an SLDS to data, we not only learn a flexible nonlinear generative model, but also learn to parse data sequences into coherent discrete units.

The generative model is as follows. At each time *t* = 1, 2, …, *T_w_*, for each worm *w*, there is a discrete latent state 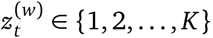. The subscript indicates the time at which this state is evaluated, and the superscript indicates which worm this state corresponds to. The states follow *Markovian* dynamics,

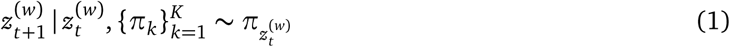

where 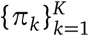 is the Markov transition matrix and *π_k_* ∈ Δ*_K_* is its *k*th row, a point in the simplex of non-negative vectors that sum to one (i.e., a probability vector). To sample the state at time *t* + 1, we draw from a distribution *π_z_t__*(w), a row of a transition matrix.

In addition to the discrete states, each worm also has a continuous latent state 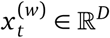, where *D* is the dimension of the continuous latent state. The continuous state captures the low-dimensional nature of neural activity and reflects correlations between neurons. The continuous states are assumed to follow conditionally affine dynamics, where the discrete state *z_t_*^(*w*)^ determines the linear dynamical system used at time *t*:

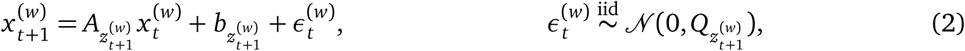

where *A_k_ ∈* ℝ*^D×D^* is a dynamics matrix, *b_k_ ∈* ℝ*^D^* is a bias vector, and 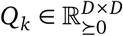 is a positive definite covariance matrix, for *k* = 1, 2, …, *K*. Again, the discrete states index into a set of linear dynamical system parameters and specify which dynamics to use when updating the continuous states.

**Table 1A:**
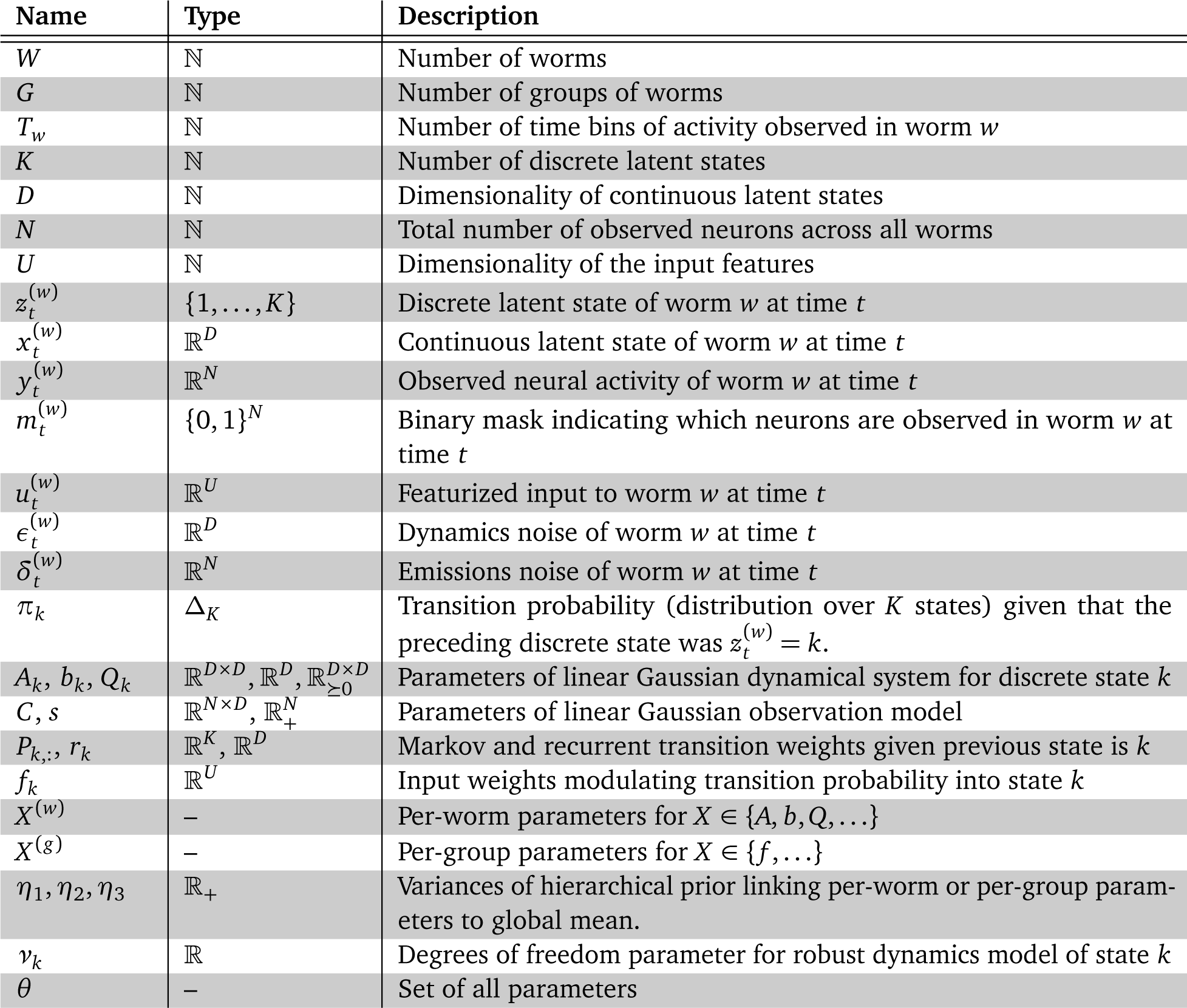
Table of generative model notation.

Finally, at each time *t* we have a linear Gaussian observation 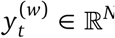, where *N* is the number of neurons. We assume that the neural activity is standardized to have mean zero and unit variance. We model the activity as a noisy observation of the underlying latent continuous state,

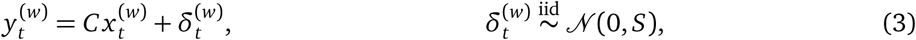

for *C ∈* ℝ*^N×D^* and *S* = diag ([*s*_1_, …, *s_N_*]) with *s_n_ >* 0. We denote the rows of *C* by vectors *c_n_*. For simplicity, we assume *C* and *S* are shared among all discrete states and all worms in our model. Overall, the system parameters consist of the discrete Markov transition matrix, the library of linear dynamical system matrices, and the neuron-specific emission parameters, which we write as

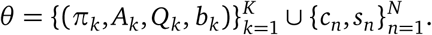

This specifies the complete data generating process for an entire population of neurons. In practice, however, we only observe a subset of this population. We indicate which entries in the data matrices are observed with a *mask* 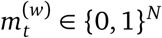, where 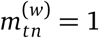 indicates that the corresponding entry 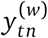 is observed. Unobserved entries, for which 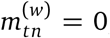, are treated as missing at random. With the diagonal covariance matrix of the observation noise, the neurons are conditionally independent given the underlying latent states, and the missing data entries effectively drop out of the joint probability density,

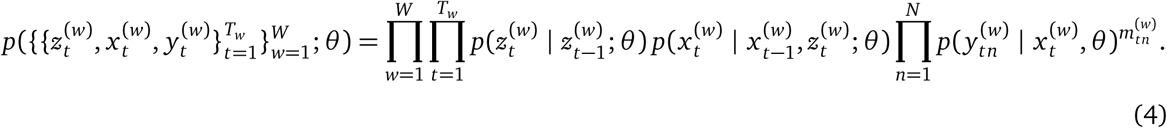

We do not specify a prior distribution on the parameters *θ* and instead estimate them via maximum likelihood.

### A.2 Extensions of the standard SLDS

To better model the *C. elegans* data, we introduce three extensions to the standard SLDS.

#### Recurrent model of discrete state transition probabilities

The first extension allows for more complex transition probabilities. We replace (1) with

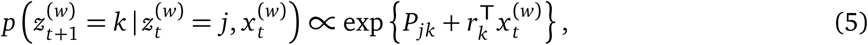

where *P ∈* ℝ*^K×K^* and *r_k_ ∈* ℝ*^D^* parameterize a map from previous discrete and continuous states to a distribution over next discrete states. The link function is the softmax function. Graphically, the recurrent model introduces an extra set of conditional dependencies in the probabilistic graphical model, as shown in Fig. D.1b. We recover the standard model when *r_k_ ≡* 0 for all *k*; in that case, *P* parameterizes an unnormalized log transition matrix.

#### Hierarchical model to allow limited variability across worms

The second extension captures shared patterns across worms while still allowing for worm-to-worm variability. We do this in two ways. First, we allow the dynamics parameters to vary slightly from worm to worm. Let (*A_k_*, *b_k_*) denote the global average affine dynamics parameters for state *k*, and let 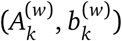 be the affine dynamics parameters specific to worm *w*. We tie the worm-specific parameters to the global parameters via a hierarchical model,

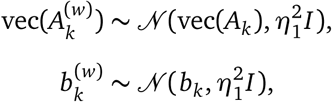

where vec(*·*) denotes the vectorization operation, which unravels a matrix into a vector. This allows worm-specific dynamics parameters 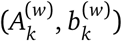 but requires that they not deviate too far from the global mean (*A_k_*, *b_k_*).

We do the same for the transition model,

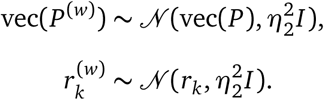

Fig. D.1c illustrates the complete probabilistic graphical model. Each worm has a set of discrete states *z*_1:*T*_, continuous states *x*_1:*T*_, and observations for a subset of neurons. The worms share a set of global dynamics parameters and emission matrices, but they also have worm-specific perturbations of the global dynamics, as well as worm-specific observation variances.

### Input-driven transition probabilities

Finally, we incorporate exogenous time-varying inputs 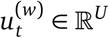 into the model by allowing them to modulate the transition probabilities. Specifically, our transition model now has the following conditionals,

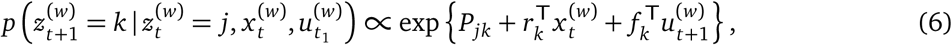

where *f_k_ ∈* ℝ*^U^* is an input weight that modulates transition probabilities as a function of the current input. Under the hierarchical model, we allow group-specific input weights 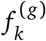 where *g* denotes the number of group index of the worm. We weakly tie the group parameters via a shared prior, 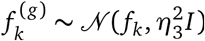.

The graphical model of the hierarchical, recurrent, input-driven SLDS is shown in Fig. D.1d.

#### Robust dynamics model

In addition to the three extensions above, we also use heavy tailed dynamics noise models. While the SLDS can capture nonlinear dynamics by composing simple linear pieces, it is still an approximation to the true data-generating process. To account for some of this model misspecification, we introduce a *robust* extension of the SLDS by allowing for heavy-tailed noise in the dynamics. Specifically, we replace (2) with

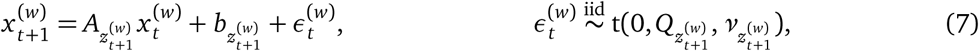

where t(*·*, *·*, *·*) denotes the multivariate-t distribution. This distribution has heavier tails than its Gaussian counterpart; it can be derived by a mixture of multivariate Gaussians with a *χ*^2^-distributed scaling of the covariance matrix. This does not incur a change in the graphical model structure.

#### B Two-step model fitting

To a first approximation, the hierarchical recurrent state space models developed here can be separated into two components: a dynamics model and an emission model. The dynamics model governs the prior distribution on the discrete and continuous states *p*(*x*, *z*; *θ*), and the emission model specifies the conditional distribution over the data given the continuous states *p*(*y | x*; *θ*). All of our models admit this factorization.

In practice, the emission model (the “likelihood”) contributes much more to the posterior distribution over the continuous states *x* than the dynamics model (the “prior”). This implies that we can get an adequate estimate of the continuous states by ignoring the prior, or by replacing it with a weak prior that admits simple posterior inference. For example, if we replace the prior on the continuous states with a simple factorized Gaussian model,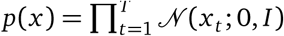, the joint distribution *p*(*x*, *y*; *θ*) is linear and Gaussian, and equivalent to linear factor analysis. The posterior mean of *x* can be found with a simple expectation maximization (EM) algorithm [Dempster et al., 1977], which accounts for missing entries in *y*. See Paninski et al. [2010] for a neuroscience-focused review of similar models and methods.

Once we have obtained a reasonable estimate of *x*, we treat it as observed and estimate the discrete states *z* and the parameters of the dynamics model using expectation maximization again. Here, the joint distribution *p*(*x*, *z*; *θ*) is equivalent to a (hierarchical, recurrent, input-driven, robust) autoregressive hidden Markov model (AR-HMM) [Hamilton, 1990, Berchtold, 1999]. Murphy [2012, Ch. 17&18] offers an overview of many state space models and extensions. The discrete states follow Markovian dynamics, as in the standard HMM, but the AR-HMM includes an extra dependency between observations. Namely, the observation *x_t_* depends on the preceding observation *x_t−_*_1_ (and possibly earlier observations as well). The EM algorithm alternates between computing the posterior distribution *p*(*z | x*; *θ*), holding the parameters *θ* fixed, and then updating the parameters to the values that maximize the expected log probability under that posterior distribution. Formally, the parameters *θ* ^(*i*)^ at iteration *i* are updated via the following procedure,

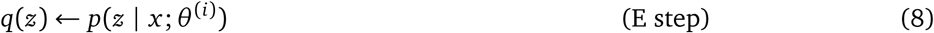

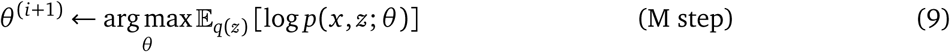

For more complex models, the M step (9) will not have a closed form solution. Instead, we use BFGS [Fletcher, 2013], a quasi-Newton optimization method, to solve for the next parameters.

Finally, it is useful to compute the most likely discrete state sequence *z^∗^* = arg max*_z_ p*(*z | x*; *θ*). This is efficiently computable via the Viterbi algorithm [Viterbi, 1967]. The code to fit these models is publicly available in the SSM package (https://github.com/slinderman/ssm).

#### C End-to-end model fitting

The two-step procedure above is fast, simple, and works well for our purposes. For completeness, however, we developed an end-to-end variational inference algorithm [Blei et al., 2017] to simultaneously learn the parameters *θ* of the generative model and infer the posterior distribution over both the discrete states *z* and continuous states *x* given these parameters and the observed data *y*. The end-to-end methods developed here yield similar results to the two-step method described above. This section is not required in order to understand the results in the main paper.

We seek parameters *ϕ* that minimize the Kullback-Leibler divergence between the approximate and true posterior, KL (*q*(*x*, *z*; *ϕ*) *I p*(*x*, *z | y*; *θ*)). In doing so, we also maximize a lower bound on the marginal log likelihood, or “evidence,” of the data since

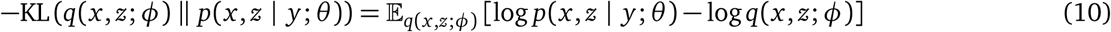

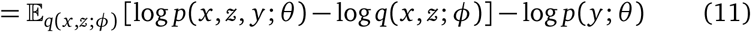

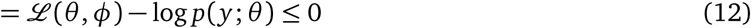

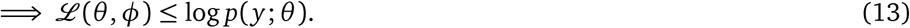

The left hand side of (13) is called the *evidence lower bound* (ELBO).

The tightness of this lower bound is determined by the quality of the posterior approximation, and the ELBO only equals the marginal log likelihood if the variational posterior equals the true posterior. The variational posterior will necessarily be approximate for nontrivial models like this, but we can still leverage the structure of the generative model to reduce this approximation error. In particular, we can compute the exact conditional distribution over the discrete states *z* given the continuous latent states *x* and use this in a structured variational approximation, *q*(*x*, *z*; *ϕ*, *θ*) = *q*(*x*; *ϕ*) *p*(*z | x*, *θ*). Substituting this approximation into the ELBO yields

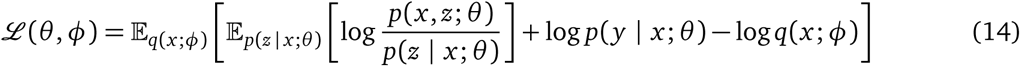

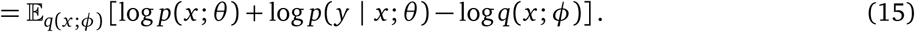

The marginal probability *p*(*x*; *θ*) is easily computable with standard HMM message passing passing routines [Rabiner, 1989].

Still, the expectation with respect to *q*(*x*; *ϕ*) is not exactly computable. Instead, we obtain an unbiased estimate of the ELBO using Monte Carlo,

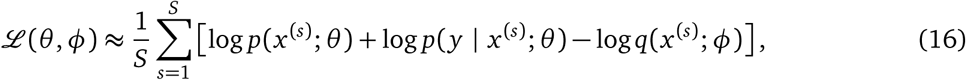

where *x*^(*s*)^ ~ *q*(*x*; ϕ).

So far we have implicitly assumed that we can efficiently sample from *q*(*x*; *ϕ*) and evaluate its density pointwise. We considered two possibilities. First, we used a mean-field variational approximation,

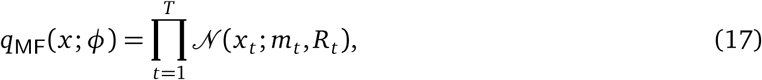

where the variational parameters are 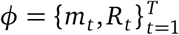. While this method is simple and easy to implement, it fails to capture temporal correlations in the continuous latent states. To address this shortcoming, we also consider a structured variational family of the form,

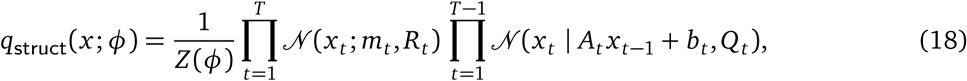

where the variational parameters are 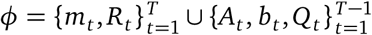. Since the dependencies are only pairwise, we can use a message passing algorithm—the Kalman filter [Kalman, 1960]—to compute the normalization constant *Z* (*ϕ*) and sample the variational posterior in *O*(*T*) time.

Alternatively, we can view *q*_struct_(*x*; *ϕ*) as a multivariate Gaussian distribution on the entire sequence of continuous latent states 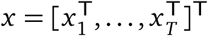 with natural parameters *J ∈* ℝ*^T D×T D^* and *h ∈* ℝ*^T D^*. *J* is the inverse covariance matrix of the multivariate Gaussian, and it has a block tridiagonal sparsity pattern, reflecting the pairwise dependencies between continuous latent states:

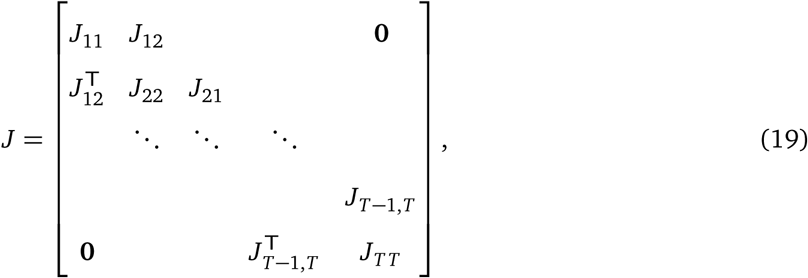

where

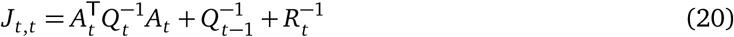

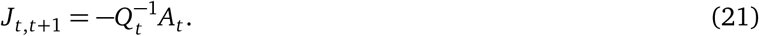

The second natural parameter determines the mean via the relation 𝔼[*x*] = *J^−^*^1^*h*. In terms of the variational parameters above, we have

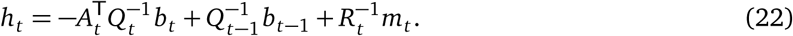

We can evaluate the density of *q*_struct_ given the Cholesky factorization *J* = *LL*^T^ using,

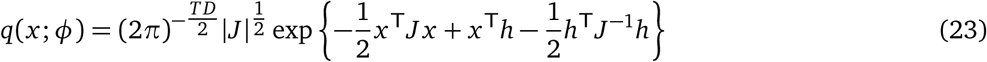

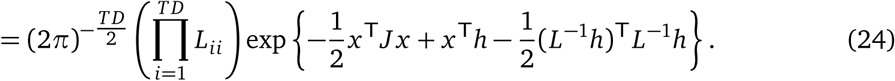

The sparsity of *J* renders the quadratic form *x* ^T^*J x* efficiently computable, and the only challenge is computing the Cholesky factorization and solving *L^−^*^1^*h*. In general, the Cholesky factorization is *O*(*T* ^3^) and the solve is *O*(*T* ^2^), but due to the sparsity of *J*, both can be computed in *O*(*T*) time. Sampling from *q*(*x*; *ϕ*) leverages the same operations. We can sample via the reparameterization,

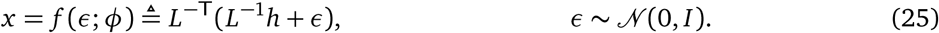

While routines for the Cholesky factorization and solves of block tridiagonal matrices are not natively available in most libraries, LAPACK does support banded matrices via `pbtrf` and `pbtrs`, and block tridiagonal matrices are a special case of banded matrices. Using banded routines incurs a small overhead but the convenience may be worthwhile. We have written C routines to convert block tridiagonal matrices to and from banded representations used by LAPACK.

The simplest method of optimizing the ELBO is via stochastic gradient ascent. We obtain an unbiased estimate of the gradient of the ELBO using the reparameterization above,

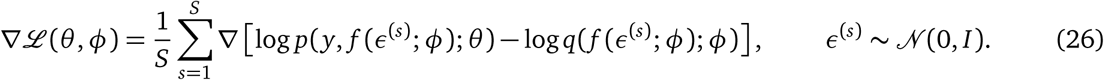

The gradient is taken with respect to both *θ* and *ϕ*.

### D Supplementary Figures

**Figure D.1:**
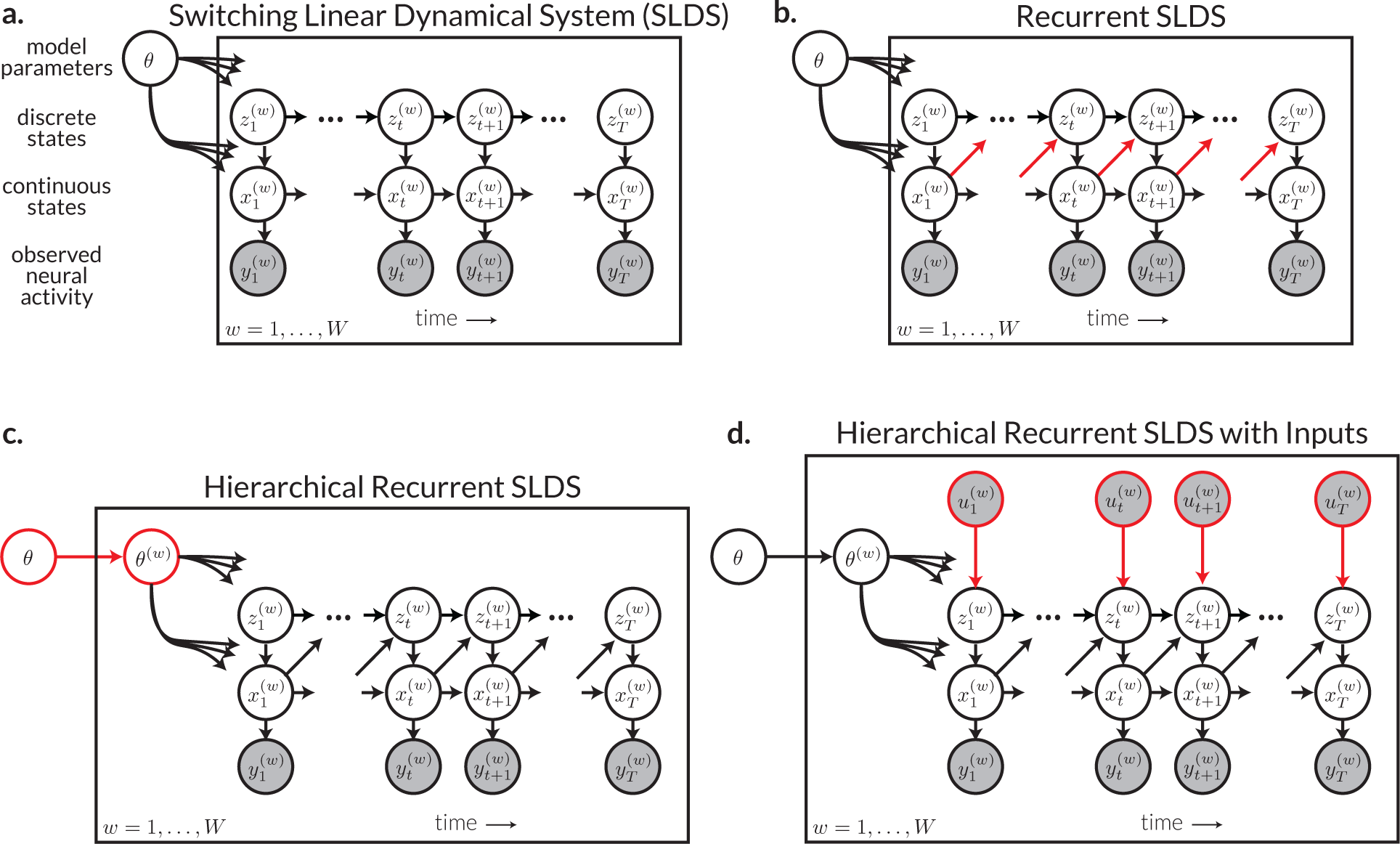
Graphical model of the standard switching linear dynamical system and its extensions. **a.** The standard SLDS has observed data variables 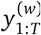 for each worm *w* 1, …, *W*. Circles denote random variables, arrows denote conditional dependencies, and filled circles indicate which variables are observed. Under this model, the observed data is a linear function of underlying continuous latent states 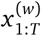 and discrete latent states 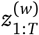. The continuous 1:*T* 1:*T* states are driven by linear dynamics that switch depending on the discrete state. The model is parameterized by transition probabilities, linear dynamics, and a mapping from continuous states to observed data, all of which are summarized in *θ*. **b.** The *Recurrent* SLDS extends the standard model with an extra set of dependencies (red arrows). Here, the discrete state transitions are determined not only by the previous discrete state, but also by the preceding continuous state. **c.** The *Hierarchical* Recurrent SLDS adds per-worm parameters *θ* ^(*w*)^, which allow for slight worm-to-worm variability. They are tied together by a prior that specifies each worm’s parameters to be centered on the global parameters *θ*. **d.** The final elaboration adds *inputs* 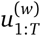 to the model. For example, we consider the oxygen concentration and change in oxygen concentration as external drivers of state transition. At the same time, we allow the parameters of these input dependencies to vary based on the genetic strain and developmental stage of the worm by including them in *θ* ^(*w*)^.

**Figure D.2:**
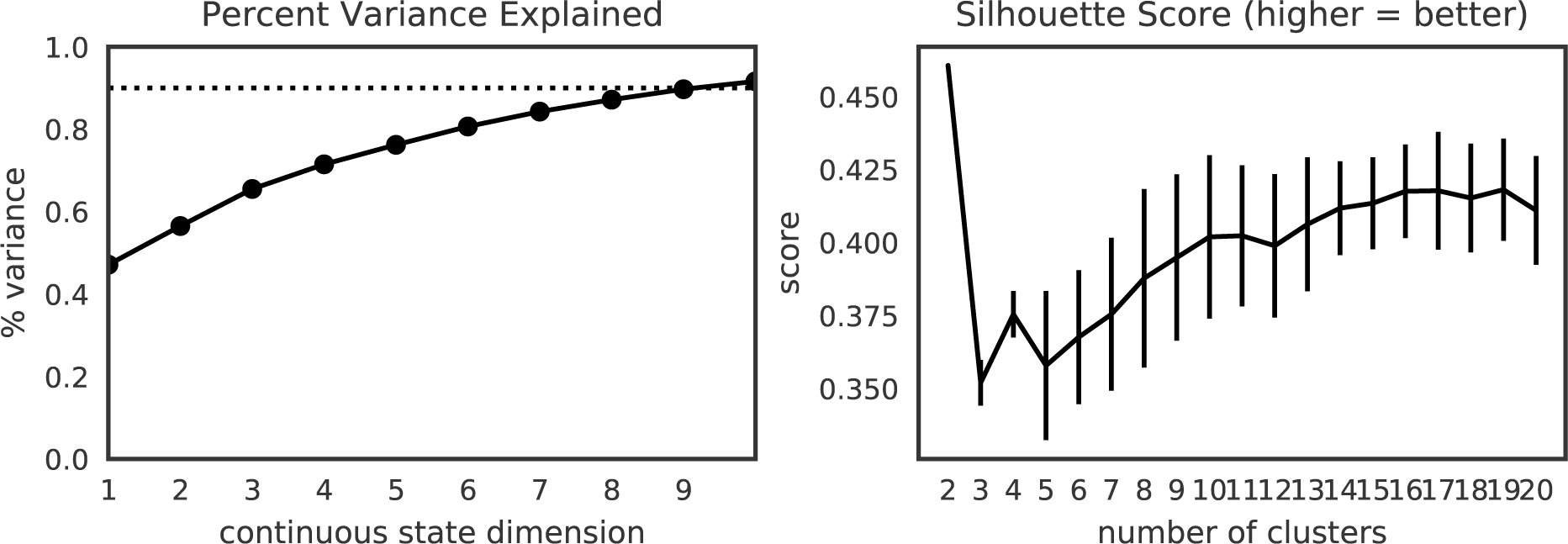
Selecting latent dimensionality and number of neuron clusters. We selected the latent dimension of *D* = 10 because it achieves *>* 90% variance explained (after reconstructing missing data). We chose the number of clusters based on where the increase in silhouette score starts to diminish. By this metric, two clusters would be ideal, but we opt for more clusters to further separate distinct functional groups of neurons.

**Figure D.3:**
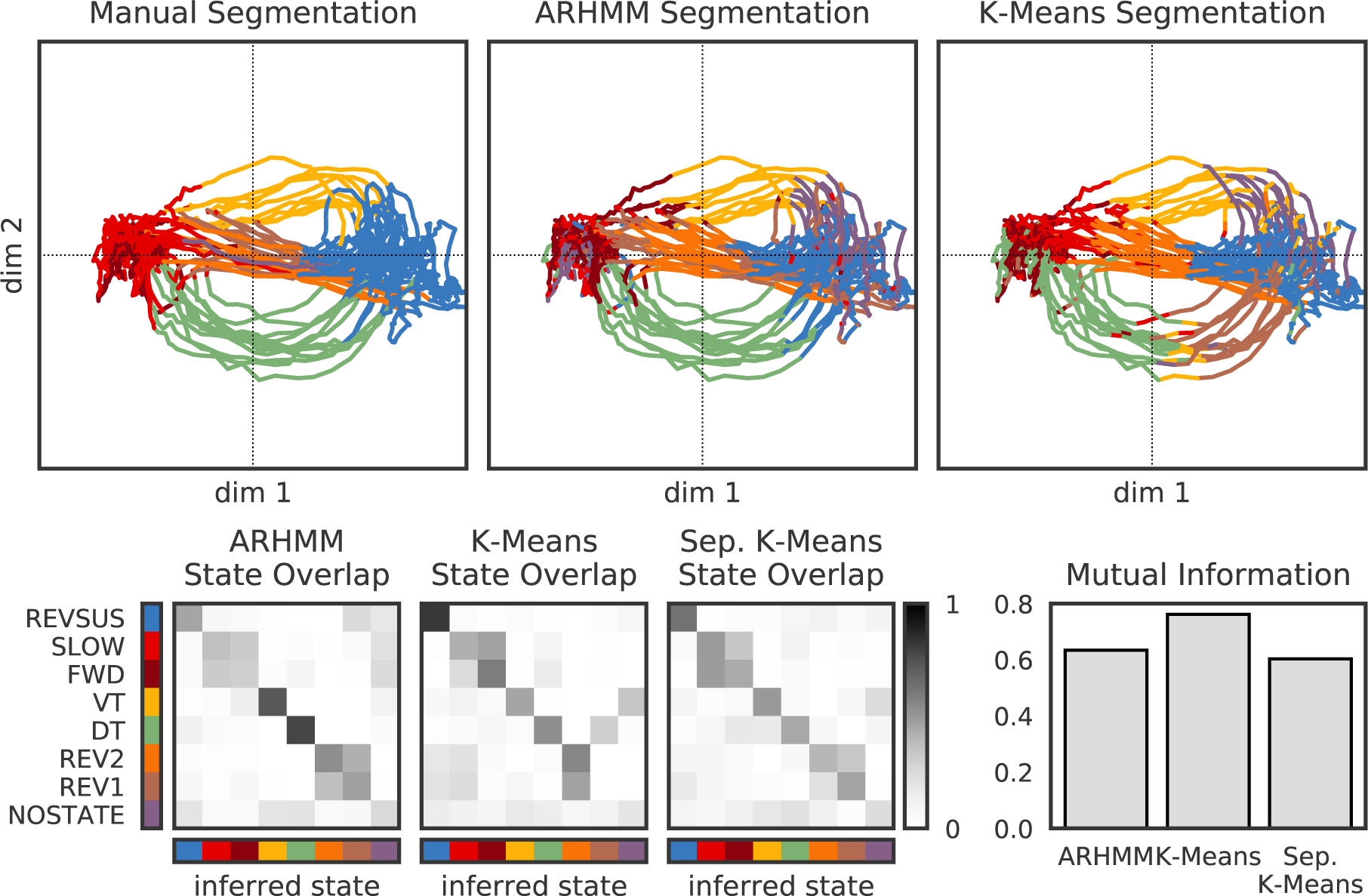
Manual segmentations can be approximately recovered without explicitly modeling dynamics, but turns are subdivided. We show the segmented trajectories under the manual discrete states, the hierarchical recurrent AR-HMM, and K-Means applied to the continuous states. Below, we show the overlap with the true states for the AR-HMM (left) compared to K-Means fit to all five worms (middle) and K-Means fit to each worm individually (right). We further quantify the overlap by calculating the mutual information between the true and inferred states under each model.

**Figure D.4:**
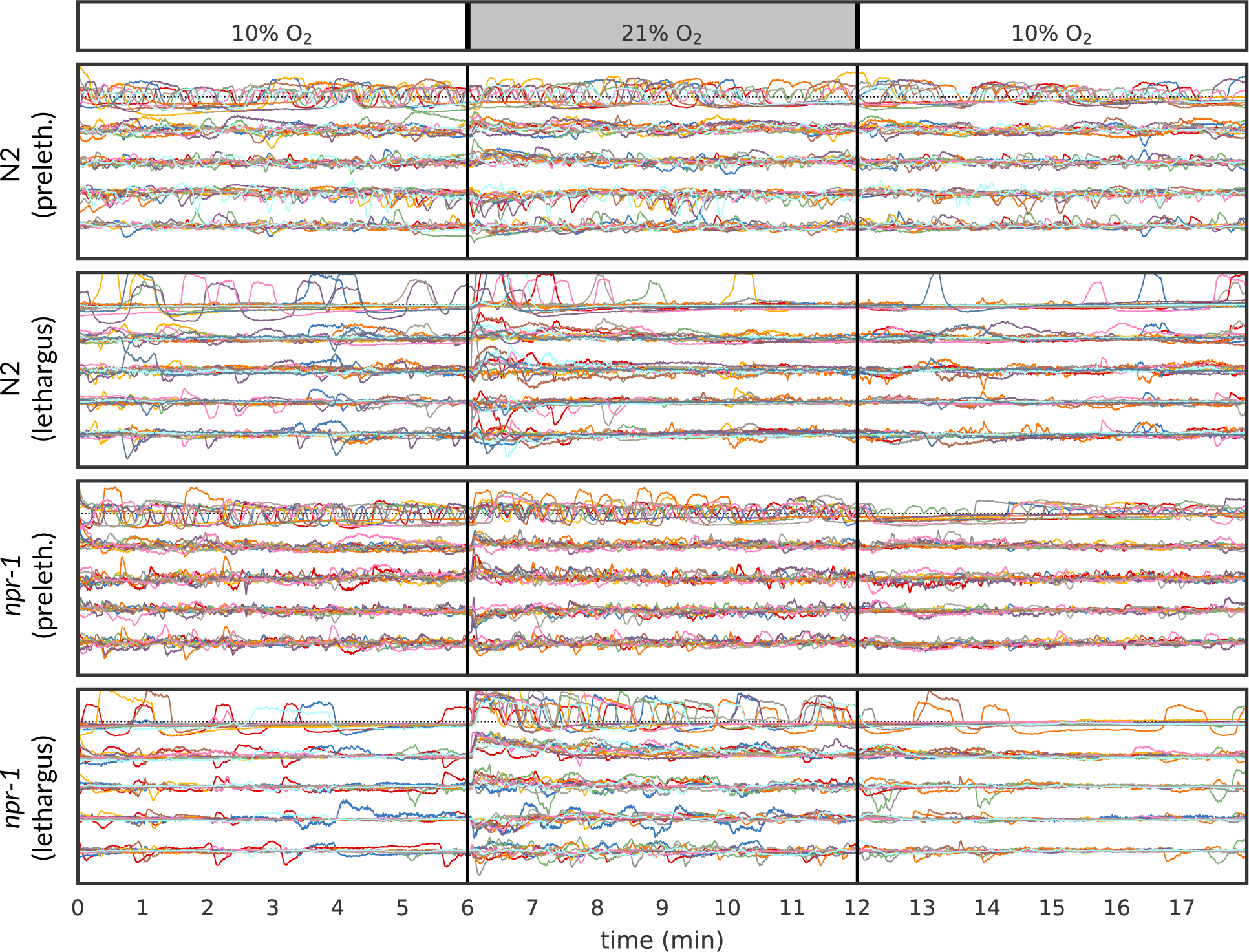
Continuous latent states for each group of worms in the Nichols et al. [2017] data. Within each group, different colors correspond to different worms. This illustrates the persistent activity of prelethargus worms throughout each 18 minute trial and the elevated activity of *npr-1* lethargus worms in the presence of increased oxygen.

**Figure D.5:**
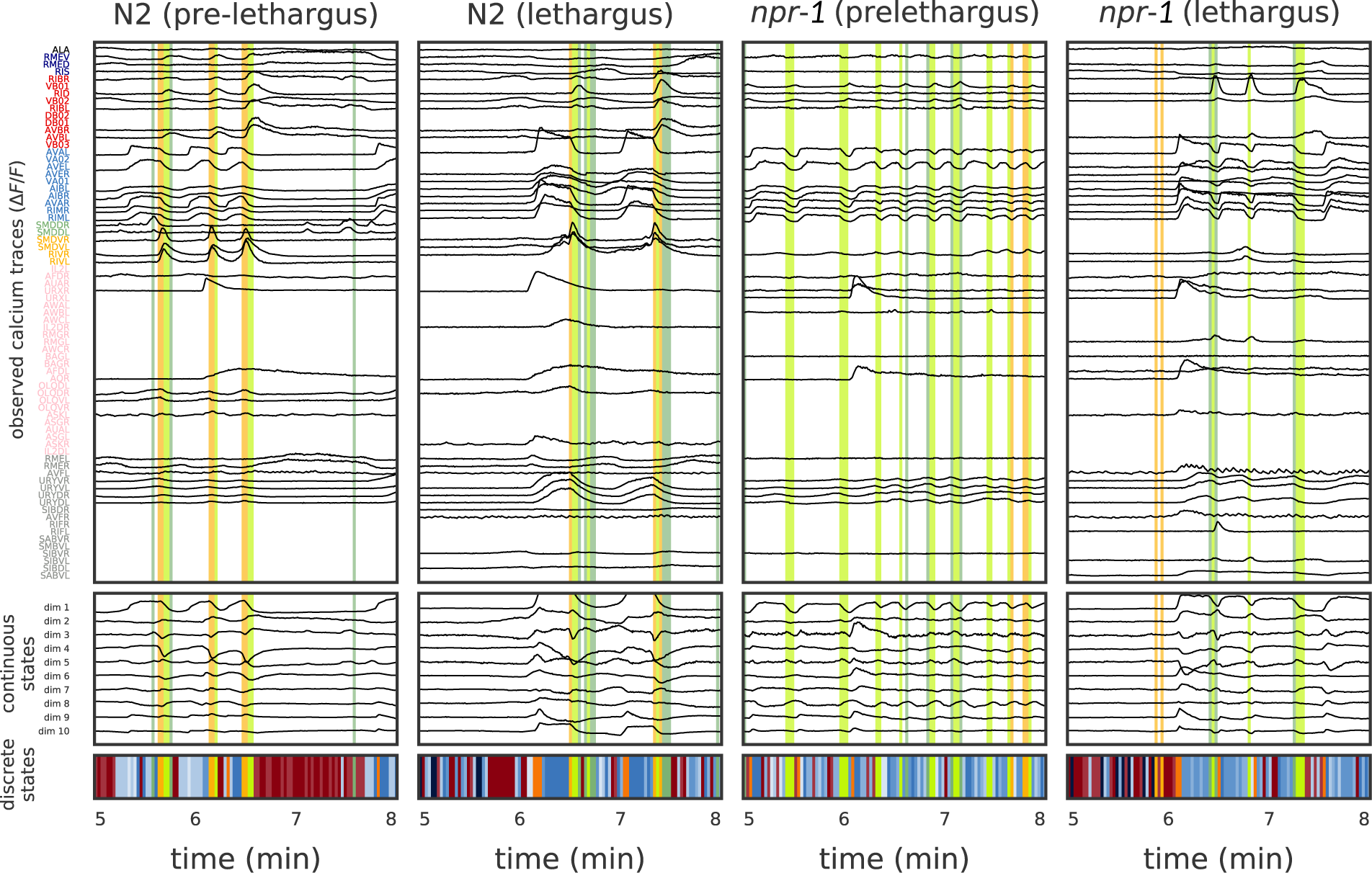
Example trajectories from each genetic strain and developmental stage in Nichols et al. [2017] data. As in Fig 1, we show the real data, the continuous states, and the inferred discrete states. We highlight reversals (orange) and turns (green/chartreuse). At 6 minutes into the recording the oxygen concentration is increased from 10% to 21%, eliciting a strong neural response in many worms.

**Figure D.6:**
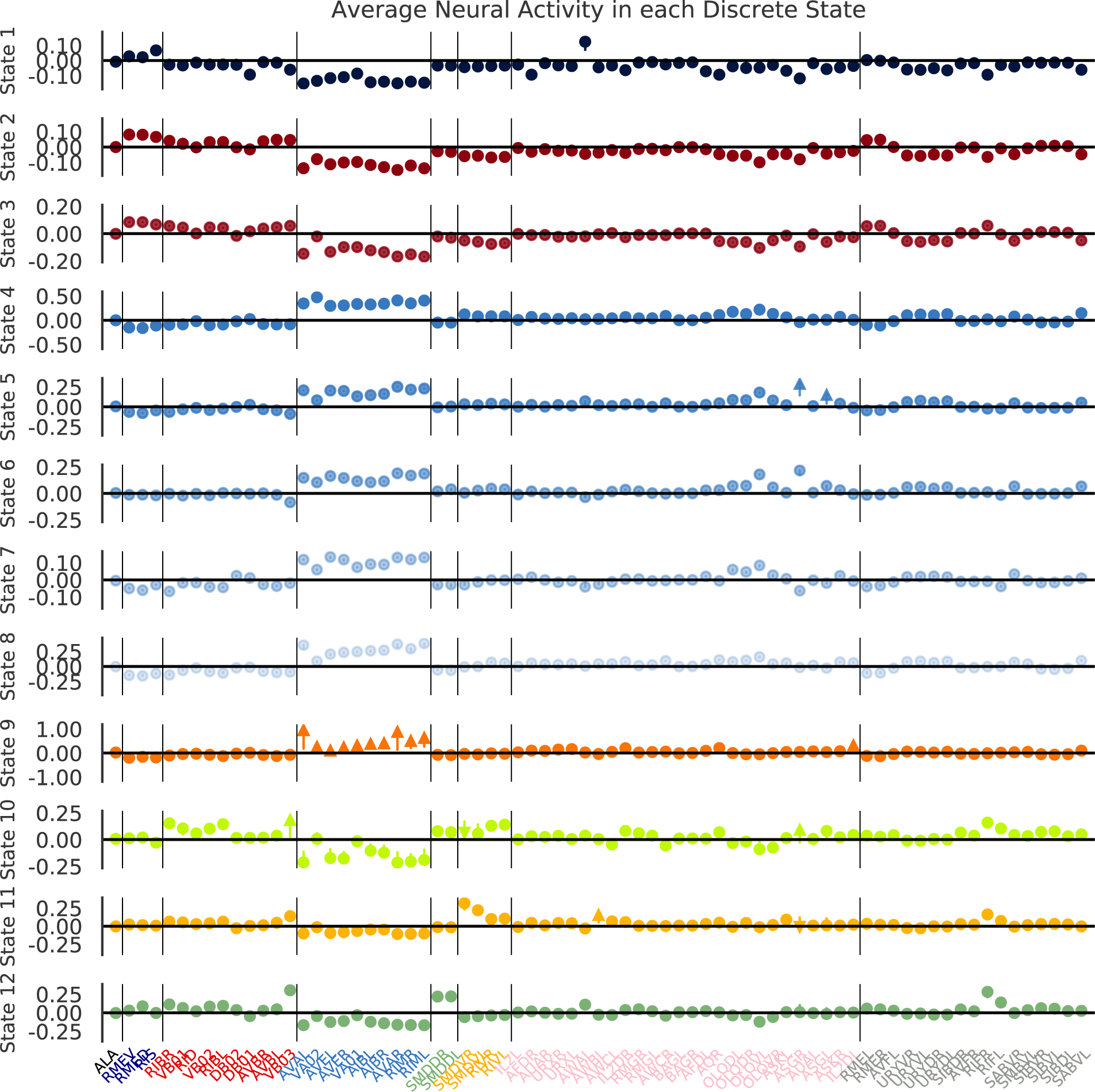
Change in neural activity for each discrete state inferred from the Nichols et al. [2017] data. This figure is analogous to Figure 4 of the main text.

### E Assessment of Neuron Functions

**Table 1E:**
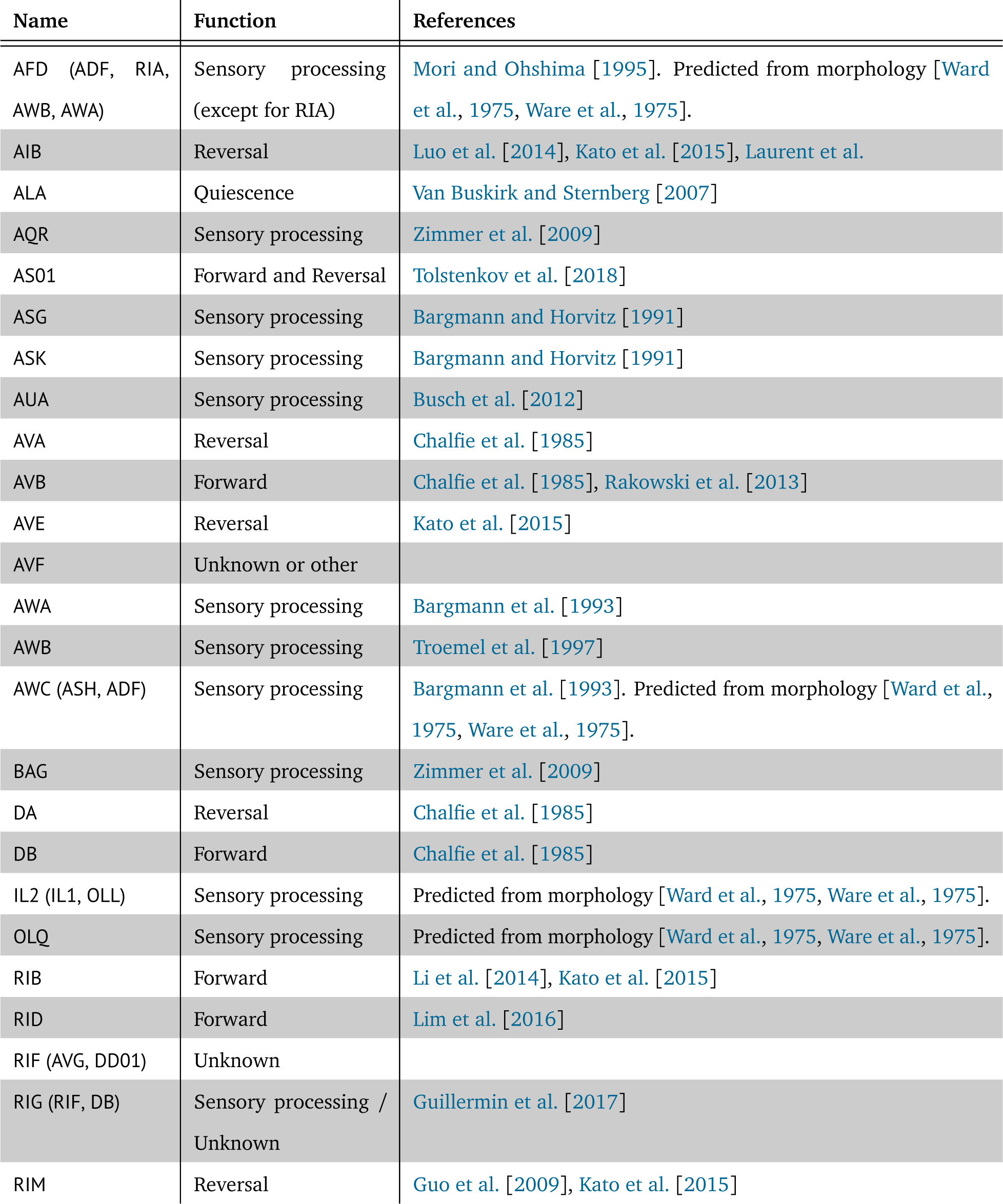
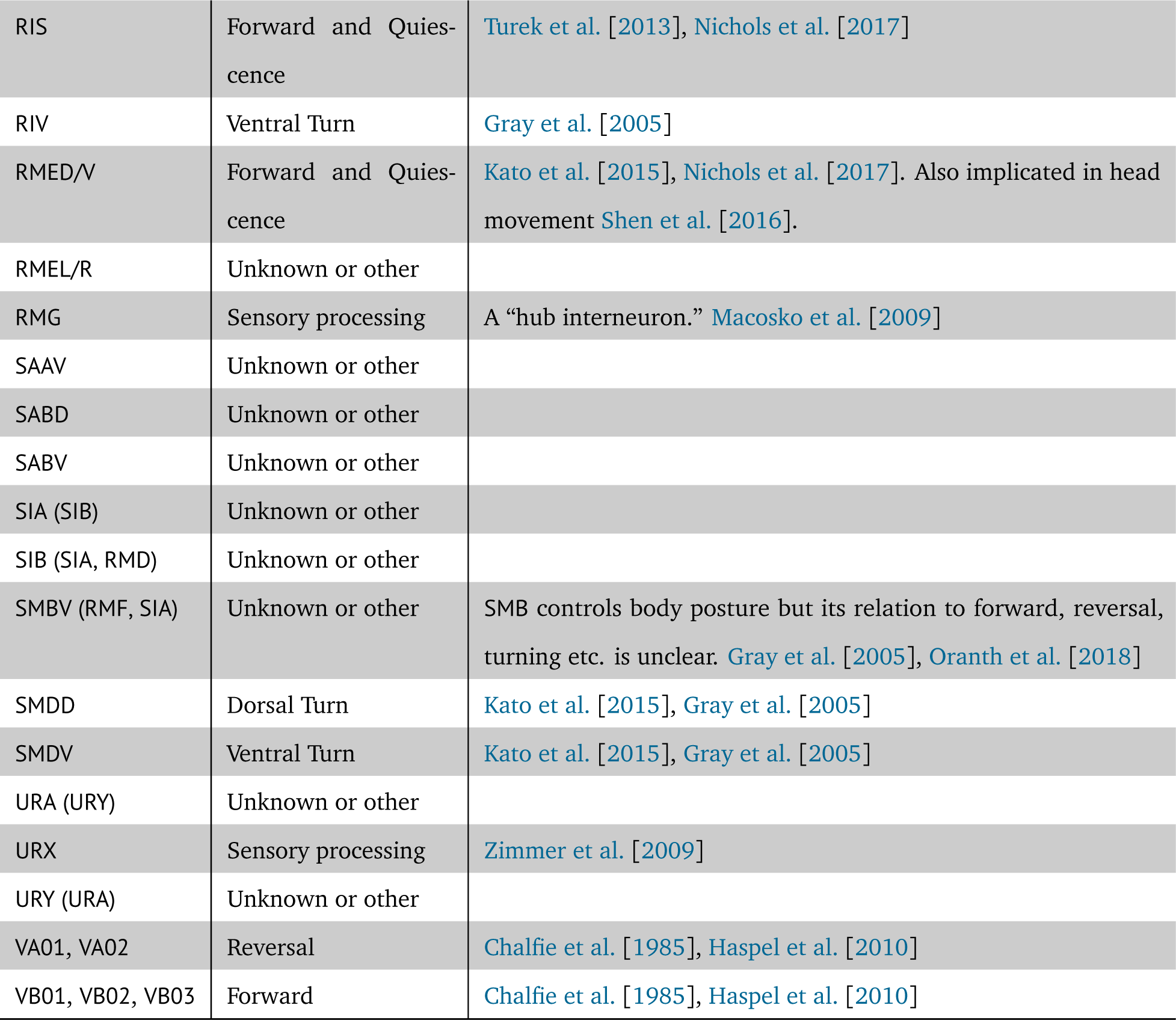
Assessment of neural function related to behaviors based on existing literature. Some neurons are difficult to identify with certainty; potential alternative identities are listed in the parentheses. The neuron functions are used for the cluster analysis in Figure 2.

## References

1. G. A. Ackerson and K.-S. Fu. On state estimation in switching environments. IEEE Transactions on Automatic Control, 15(1):10–17, 1970.

2. Z. Altun, L. Herndon, C. Wolkow, C. Crocker, R. Lints, and D.e. Hall. Wormatlas, 2018. URL http://www.wormatlas.org.

3. M. Aoi and J. W. Pillow. Model-based targeted dimensionality reduction for neuronal population data. In Advances in Neural Information Processing Systems, pages 6689–6698, 2018.

4. Y. Bar-Shalom and X.-R. Li. Estimation and tracking. Artech House, Boston, MA, 1993.

5. C. I. Bargmann and H. R. Horvitz. Chemosensory neurons with overlapping functions direct chemotaxis to multiple chemicals in C. elegans. Neuron, 7(5):729–742, 1991.

6. C. I. Bargmann, E. Hartwieg, and H. R. Horvitz. Odorant-selective genes and neurons mediate olfaction in C. elegans. Cell, 74(3):515–527, 1993.

7. C. J. Bartholomew, M. Knott, and I. Moustaki. Latent variable models and factor analysis: A unified approach, volume 904. John Wiley & Sons, 2011.

8. C. Batty, J. Merel, N. Brackbill, A. Heitman, A. Sher, A. Litke, E. J. Chichilnisky, and L. Paninski. Multilayer recurrent network models of primate retinal ganglion cell responses. International Conference on Learning Representations, 2017.

9. S. Behseta, R. E. Kass, and G. L. Wallstrom. Hierarchical models for assessing variability among functions. Biometrika, 92(2):419–434, 2005.

10. Y. Bengio and P. Frasconi. An input-output HMM architecture. In Advances in Neural Information Processing Systems, pages 427–434, 1995.

11. A. Berchtold. The double chain Markov model. Communications in Statistics-Theory and Methods, 28 (11):2569–2589, 1999.

12. D. M. Blei, A. Kucukelbir, and J. D. McAuliffe. Variational inference: A review for statisticians. Journal of the American Statistical Association, 112(518):859–877, 2017.

13. C. Brennan and A. Proekt. Universality of macroscopic neuronal dynamics in Caenorhabditis elegans. arXiv preprint arXiv:1711.08533, 2017.

14. K. E. Busch, P. Laurent, Z. Soltesz, R. J. Murphy, O. Faivre, B. Hedwig, M. Thomas, H. L. Smith, and M. De Bono. Tonic signaling from O*_2_* sensors sets neural circuit activity and behavioral state. Nature neuroscience, 15(4):581, 2012.

15. M. Chalfie, J. E. Sulston, J. G. White, E. Southgate, J. N. Thomson, and S. Brenner. The neural circuit for touch sensitivity in Caenorhabditis elegans. Journal of Neuroscience, 5(4):956–964, 1985.

16. C.-B. Chang and M. Athans. State estimation for discrete systems with switching parameters. *IEEE Transactions on Aerospace and Electronic Systems*, AES-14(3):418–425, 1978.

17. Z. Chen, S. N. Gomperts, J. Yamamoto, and M. A. Wilson. Neural representation of spatial topology in the rodent hippocampus. Neural Computation, 26(1):1–39, 2014.

18. M. M. Churchland, J. P. Cunningham, M. T. Kaufman, J. D. Foster, P. Nuyujukian, S. I. Ryu, and K. V. Shenoy. Neural population dynamics during reaching. Nature, 487(7405):51, 2012.

19. A. C. Costa, T. Ahamed, and G. J. Stephens. Adaptive, locally linear models of complex dynamics. Proceedings of the National Academy of Sciences, 116(5):1501–1510, 2019.

20. A. Cronin, I. H. Stevenson, M. Sur, and K. P. Körding. Hierarchical Bayesian modeling and Markov chain Monte Carlo sampling for tuning-curve analysis. J. Neurophysiol., 103(1):591–602, Jan. 2010.

21. J. P. Cunningham and M. B. Yu. Dimensionality reduction for large-scale neural recordings. Nature neuroscience, 17(11):1500, 2014.

22. P. Dayan and L. F. Abbott. Theoretical neuroscience: computational and mathematical modeling of neural systems. MIT press, 2001.

23. A. P. Dempster, N. M. Laird, and D. B. Rubin. Maximum likelihood from incomplete data via the EM algorithm. Journal of the Royal Statistical Society: Series B (Methodological), 39(1):1–22, 1977.

24. L. Duncker and M. Sahani. Temporal alignment and latent Gaussian process factor inference in population spike trains. In Advances in Neural Information Processing Systems, pages 10466–10476, 2018.

25. R. Fletcher. Practical methods of optimization. John Wiley & Sons, 2013.

26. E. Fox, E. B. Sudderth, M. I. Jordan, and A. S. Willsky. Nonparametric Bayesian learning of switching linear dynamical systems. Advances in Neural Information Processing Systems, pages 457–464, 2009.

27. Y. Gao, E. W. Archer, L. Paninski, and J. P. Cunningham. Linear dynamical neural population models through nonlinear embeddings. In Advances in Neural Information Processing Systems, pages 163–171, 2016.

28. A. Gelman and J. Hill. Data analysis using regression and multilevel/hierarchical models. Cambridge University Press, 2006.

29. Z. Ghahramani and G. E. Hinton. Switching state-space models. Technical report, University of Toronto, 1996.

30. A. Gordus, N. Pokala, S. Levy, S. W. Flavell, and C. I. Bargmann. Feedback from network states generates variability in a probabilistic olfactory circuit. Cell, 161(2):215–227, 2015.

31. J. M. Gray, D. S. Karow, H. Lu, A. J. Chang, J. S. Chang, R. E. Ellis, M. A. Marletta, and C. I. Bargmann. Oxygen sensation and social feeding mediated by a C. elegans guanylate cyclase homologue. Nature, 430(6997):317, 2004.

32. J. M. Gray, J. J. Hill, and C. I. Bargmann. A circuit for navigation in Caenorhabditis elegans. Proceedings of the National Academy of Sciences, 102(9):3184–3191, 2005.

33. M. L. Guillermin, M. A. Carrillo, and E. A. Hallem. A single set of interneurons drives opposite behaviors in C. elegans. Current Biology, 27(17):2630–2639, 2017.

34. Z. V. Guo, A. C. Hart, and S. Ramanathan. Optical interrogation of neural circuits in Caenorhabditis elegans. Nature Methods, 6(12):891, 2009.

35. J. D. Hamilton. Analysis of time series subject to changes in regime. Journal of econometrics, 45(1): 39–70, 1990.

36. P. J. Harrison and C. F. Stevens. Bayesian forecasting. Journal of the Royal Statistical Society. Series B (Methodological), pages 205–247, 1976.

37. G. Haspel, M. J. O’Donovan, and A. C. Hart. Motoneurons dedicated to either forward or backward locomotion in the nematode Caenorhabditis elegans. Journal of Neuroscience, 30(33):11151–11156, 2010.

38. D. Hernandez, A. K. Moretti, Z. Wei, S. Saxena, J. Cunningham, and L. Paninski. A novel variational family for hidden nonlinear Markov models. arXiv preprint arXiv:1811.02459, 2018.

39. M. J. Johnson and A. S. Willsky. Bayesian nonparametric hidden semi-Markov models. Journal of Machine Learning Research, 14(Feb):673–701, 2013.

40. L. M. Jones, A. Fontanini, B. F. Sadacca, P. Miller, and D. B. Katz. Natural stimuli evoke dynamic sequences of states in sensory cortical ensembles. Proceedings of the National Academy of Sciences, 104(47): 18772–18777, 2007.

41. R. E. Kalman. A new approach to linear filtering and prediction problems. Journal of Basic Engineering, 82(1):35–45, 1960.

42. R. Kato, H. S. Kaplan, T. Schrödel, S. Skora, T. H. Lindsay, E. Yemini, S. Lockery, and M. Zimmer. Global brain dynamics embed the motor command sequence of Caenorhabditis elegans. Cell, 163(3):656–669, 2015.

43. R. Kawano, M. D. Po, S. Gao, G. Leung, W. S. Ryu, and M. Zhen. An imbalancing act: gap junctions reduce the backward motor circuit activity to bias C. elegans for forward locomotion. Neuron, 72(4): 572–586, 2011.

44. D. Kobak, W. Brendel, C. Constantinidis, C. E. Feierstein, A. Kepecs, Z. F. Mainen, X.-L. Qi, R. Romo, N. Uchida, and C. K. Machens. Demixed principal component analysis of neural population data. eLife, 5:e10989, 2016.

45. A. Kocabas, C.-H. Shen, Z. V. Guo, and S. Ramanathan. Controlling interneuron activity in Caenorhabditis elegans to evoke chemotactic behaviour. Nature, 490(7419):273, 2012.

46. P. Laurent, Z. Soltesz, G. M. Nelson, C. Chen, F. Arellano-Carbajal, E. Levy, and M. de Bono. Decoding a neural circuit controlling global animal state in C. elegans.

47. J. J. Li, H. Huang, P. J. Bickel, and S. E. Brenner. Comparison of D. melanogaster and C. elegans developmental stages, tissues, and cells by modENCODE RNA-seq data. Genome research, 24(7): 1086–1101, 2014.

48. M. A. Lim, J. Chitturi, V. Laskova, J. Meng, D. Findeis, A. Wiekenberg, B. Mulcahy, L. Luo, Y. Li, Y. Lu, et al. Neuroendocrine modulation sustains the C. elegans forward motor state. eLife, 5:e19887, 2016.

49. S. W. Linderman, M. J. Johnson, M. A. Wilson, and Z. Chen. A Bayesian nonparametric approach for uncovering rat hippocampal population codes during spatial navigation. Journal of Neuroscience Methods, 263:36–47, 2016.

50. S. W. Linderman, M. J. Johnson, A. C. Miller, R. P. Adams, D. M. Blei, and L. Paninski. Bayesian learning and inference in recurrent switching linear dynamical systems. In Proceedings of the 20th International Conference on Artificial Intelligence and Statistics (AISTATS), 2017.

51. S. W. Linderman, G. E. Mena, H. Cooper, L. Paninski, and J. P. Cunningham. Reparameterizing the Birkhoff polytope for variational permutation inference. In Proceedings of the 21st International Conference on Artificial Intelligence and Statistics (AISTATS), 2018.

52. M. Liu, A. K. Sharma, J. W. Shaevitz, and A. M. Leifer. Temporal processing and context dependency in Caenorhabditis elegans response to mechanosensation. eLife, 7:e36419, 2018.

53. L. Luo, Q. Wen, J. Ren, M. Hendricks, M. Gershow, Y. Qin, J. Greenwood, E. R. Soucy, M. Klein, H. K. Smith-Parker, et al. Dynamic encoding of perception, memory, and movement in a C. elegans chemotaxis circuit. Neuron, 82(5):1115–1128, 2014.

54. J. H. Macke, L. Buesing, J. P. Cunningham, M. Y. Byron, K. V. Shenoy, and M. Sahani. Empirical models of spiking in neural populations. In Advances in Neural Information Processing Systems, pages 1350–1358, 2011.

55. E. Z. Macosko, N. Pokala, E. H. Feinberg, S. H. Chalasani, R. A. Butcher, J. Clardy, and C. I. Bargmann. A hub-and-spoke circuit drives pheromone attraction and social behaviour in C. elegans. Nature, 458 (7242):1171, 2009.

56. G. Mena, D. Belanger, S. Linderman, and J. Snoek. Learning latent permutations with Gumbel-Sinkhorn networks. International Conference on Learning Representations, 2018.

57. I. Mori and Y. Ohshima. Neural regulation of thermotaxis in Caenorhabditis elegans. Nature, 376(6538): 344, 1995.

58. K. P. Murphy. Switching Kalman filters. Technical report, Compaq Cambridge Research, 1998.

59. K. P. Murphy. Hidden semi-Markov models (HSMMs). Technical report, MIT, 2002.

60. K. P. Murphy. Machine Learning: A Probabilistic Perspective. MIT press, 2012.

61. J. Nassar, S. Linderman, M. Bugallo, and I. M. Park. Tree-structured recurrent switching linear dynamical systems for multi-scale modeling. In International Conference on Learning Representations, 2019.

62. J. P. Nguyen, F. B. Shipley, A. N. Linder, G. S. Plummer, M. Liu, S. U. Setru, J. W. Shaevitz, and A. M. Leifer. Whole-brain calcium imaging with cellular resolution in freely behaving Caenorhabditis elegans. Proceedings of the National Academy of Sciences, 113(8):E1074–E1081, 2016.

63. A. L. Nichols, T. Eichler, R. Latham, and M. Zimmer. A global brain state underlies C. elegans sleep behavior. Science, 356(6344):eaam6851, 2017.

64. M. Nonnenmacher, S. C. Turaga, and J. H. Macke. Extracting low-dimensional dynamics from multiple large-scale neural population recordings by learning to predict correlations. In Advances in Neural Information Processing Systems, pages 5702–5712, 2017.

65. A. Oranth, C. Schultheis, O. Tolstenkov, K. Erbguth, J. Nagpal, D. Hain, M. Brauner, S. Wabnig, W. S. Costa, R. D. McWhirter, et al. Food sensation modulates locomotion by dopamine and neuropeptide signaling in a distributed neuronal network. Neuron, 100(6):1414–1428, 2018.

66. C. Pandarinath, D. J. O’Shea, J. Collins, R. Jozefowicz, S. D. Stavisky, J. C. Kao, E. M. Trautmann, M. T. Kaufman, S. I. Ryu, L. R. Hochberg, et al. Inferring single-trial neural population dynamics using sequential auto-encoders. Nature Methods, 2018.

67. L. Paninski, Y. Ahmadian, D. G. Ferreira, S. Koyama, K. R. Rad, M. Vidne, J. Vogelstein, and W. Wu. A new look at state-space models for neural data. Journal of Computational Neuroscience, 29(1-2):107–126, 2010.

68. O. Perez, R. E. Kass, and H. Merchant. Trial time warping to discriminate stimulus-related from movement-related neural activity. Journal of Neuroscience Methods, 212(2):203–210, 2013.

69. B. Petreska, M. Y. Byron, J. P. Cunningham, G. Santhanam, S. I. Ryu, K. V. Shenoy, and M. Sahani. Dynamical segmentation of single trials from population neural data. In Advances in Neural Information Processing Systems, pages 756–764, 2011.

70. D. Pfau, E. A. Pnevmatikakis, and L. Paninski. Robust learning of low-dimensional dynamics from large neural ensembles. In Advances in Neural Information Processing Systems, pages 2391–2399, 2013.

71. B. J. Piggott, J. Liu, Z. Feng, S. A. Wescott, and X. S. Xu. The neural circuits and synaptic mechanisms underlying motor initiation in C. elegans. Cell, 147(4):922–933, 2011.

72. R. Prevedel, Y.-G. Yoon, M. Hoffmann, N. Pak, G. Wetzstein, S. Kato, T. Schrödel, R. Raskar, M. Zimmer, E. S. Boyden, et al. Simultaneous whole-animal 3d imaging of neuronal activity using light-field microscopy. Nature Methods, 11(7):727–730, 2014.

73. L. R. Rabiner. A tutorial on hidden Markov models and selected applications in speech recognition. Proceedings of the IEEE, 77(2):257–286, 1989.

74. D. M. Raizen, J. E. Zimmerman, M. H. Maycock, U. D. Ta, Y.-j. You, M. V. Sundaram, and A. I. Pack. Lethargus is a Caenorhabditis elegans sleep-like state. Nature, 451(7178):569, 2008.

75. F. Rakowski, J. Srinivasan, P. W. Sternberg, and J. Karbowski. Synaptic polarity of the interneuron circuit controlling C. elegans locomotion. Frontiers in Computational Neuroscience, 7:128, 2013.

76. M. Scholz, A. N. Linder, F. Randi, A. K. Sharma, X. Yu, J. W. Shaevitz, and A. Leifer. Predicting natural behavior from whole-brain neural dynamics. bioRxiv, 2018. doi: 10.1101/445643.

77. T. Schrödel, R. Prevedel, K. Aumayr, M. Zimmer, and A. Vaziri. Brain-wide 3d imaging of neuronal activity in Caenorhabditis elegans with sculpted light. Nature Methods, 10(10):1013–1020, 2013.

78. Y. Shen, Q. Wen, H. Liu, C. Zhong, Y. Qin, G. Harris, T. Kawano, M. Wu, T. Xu, A. D. Samuel, et al. An extrasynaptic GABAergic signal modulates a pattern of forward movement in Caenorhabditis elegans. eLife, 5:e14197, 2016.

79. A. C. Smith and E. N. Brown. Estimating a state-space model from point process observations. Neural Computation, 15(5):965–991, 2003.

80. D. Soudry, S. Keshri, P. Stinson, M.-h. Oh, G. Iyengar, and L. Paninski. Efficient “shotgun” inference of neural connectivity from highly sub-sampled activity data. PLoS Computational Biology, 11(10): e1004464, 2015.

81. J. Taghia, W. Cai, S. Ryali, J. Kochalka, J. Nicholas, T. Chen, and V. Menon. Uncovering hidden brain state dynamics that regulate performance and decision-making during cognition. Nature Communications, 9 (1):2505, 2018.

82. O. Tolstenkov, P. Van der Auwera, W. S. Costa, O. Bazhanova, T. M. Gemeinhardt, A. C. Bergs, and A. Gottschalk. Functionally asymmetric motor neurons contribute to coordinating locomotion of Caenorhabditis elegans. eLife, 7:e34997, 2018.

83. E. R. Troemel, B. E. Kimmel, and C. I. Bargmann. Reprogramming chemotaxis responses: sensory neurons define olfactory preferences in C. elegans. Cell, 91(2):161–169, 1997.

84. S. Turaga, L. Buesing, A. M. Packer, H. Dalgleish, N. Pettit, M. Hausser, and J. H. Macke. Inferring neural population dynamics from multiple partial recordings of the same neural circuit. In Advances in Neural Information Processing Systems, pages 539–547, 2013.

85. M. Turek, I. Lewandrowski, and H. Bringmann. An AP2 transcription factor is required for a sleep-active neuron to induce sleep-like quiescence in C. elegans. Current Biology, 23(22):2215–2223, 2013.

86. C. Van Buskirk and P. W. Sternberg. Epidermal growth factor signaling induces behavioral quiescence in Caenorhabditis elegans. Nature neuroscience, 10(10):1300, 2007.

87. V. Venkatachalam, N. Ji, X. Wang, C. Clark, J. K. Mitchell, M. Klein, C. J. Tabone, J. Florman, H. Ji, J. Greenwood, A. D. Chisholm, J. Srinivasan, M. Alkema, M. Zhen, and A. D. T. Samuel. Pan-neuronal imaging in roaming Caenorhabditis elegans. Proceedings of the National Academy of Sciences, 2015. ISSN 0027-8424. doi: 10.1073/pnas.1507109113.

88. A. Viterbi. Error bounds for convolutional codes and an asymptotically optimum decoding algorithm. IEEE transactions on Information Theory, 13(2):260–269, 1967.

89. S. Ward, N. Thomson, J. G. White, and S. Brenner. Electron microscopical reconstruction of the anterior sensory anatomy of the nematode Caenorhabditis elegans. Journal of Comparative Neurology, 160(3): 313–337, 1975.

90. R. W. Ware, D. Clark, K. Crossland, and R. L. Russell. The nerve ring of the nematode Caenorhabditis elegans: sensory input and motor output. Journal of Comparative Neurology, 162(1):71–110, 1975.

91. Z. Wei, H. Inagaki, N. Li, K. Svoboda, and S. Druckmann. An orderly single-trial organization of population dynamics in premotor cortex predicts behavioral variability. bioRxiv, page 376830, 2018.

92. M. R. Whiteway and D. A. Butts. Revealing unobserved factors underlying cortical activity with a rectified latent variable model applied to neural population recordings. Journal of neurophysiology, 117(3): 919–936, 2016.

93. A. Wu, N. G. Roy, S. Keeley, and J. W. Pillow. Gaussian process based nonlinear latent structure discovery in multivariate spike train data. In Advances in Neural Information Processing Systems, pages 3496–3505, 2017.

94. E. Yemini. Fast whole-brain imaging with complete neural identity in C. elegans. In Connectome to behaviour: modelling C. elegans at cellular resolution. The Royal Society, 2018. URL https://royalsociety.org/science-events-and-lectures/2018/01/mind-of-a-worm/.

95. B. M. Yu, J. P. Cunningham, G. Santhanam, S. I. Ryu, K. V. Shenoy, and M. Sahani. Gaussian-process factor analysis for low-dimensional single-trial analysis of neural population activity. In Advances in Neural Information Processing Systems, pages 1881–1888, 2009.

96. S.-Z. Yu. Hidden semi-Markov models. Artificial intelligence, 174(2):215–243, 2010.

97. Y. Zhao and I. M. Park. Variational latent Gaussian process for recovering single-trial dynamics from population spike trains. Neural Computation, 29(5):1293–1316, 2017.

98. M. Zimmer, J. M. Gray, N. Pokala, A. J. Chang, D. S. Karow, M. A. Marletta, M. L. Hudson, D. B. Morton, N. Chronis, and C. I. Bargmann. Neurons detect increases and decreases in oxygen levels using distinct guanylate cyclases. Neuron, 61(6):865–879, 2009.

99. W. Zou, A. Shen, X. Dong, M. Tugizova, Y. K. Xiang, and K. Shen. A multi-protein receptor-ligand complex underlies combinatorial dendrite guidance choices in C. elegans. eLife, 5:e18345, 2016.

